# Centrosome amplification primes for apoptosis and favors the response to chemotherapy in ovarian cancer beyond multipolar divisions

**DOI:** 10.1101/2023.07.28.550973

**Authors:** Frances Edwards, Giulia Fantozzi, Anthony Y. Simon, Jean-Philippe Morretton, Aurelie Herbette, Andrea E. Tijhuis, Rene Wardenaar, Stacy Foulane, Simon Gemble, Diana C.J. Spierings, Floris Foijer, Odette Mariani, Anne Vincent-Salomon, Sergio Roman Roman, Xavier Sastre-Garau, Oumou Goundiam, Renata Basto

## Abstract

Centrosome amplification is a feature of cancer cells associated with chromosome instability and invasiveness. Enhancing chromosome instability and subsequent cancer cell death via centrosome unclustering and multipolar divisions is an aimed-for therapeutic approach. Here we show that centrosome amplification favors responses to conventional chemotherapy independently of multipolar divisions and chromosome instability. We perform single-cell live imaging of chemotherapy responses in epithelial ovarian cancer cell lines and observe increased cell death when centrosome amplification is induced. By correlating cell fate with mitotic behaviors, we show that enhanced cell death occurs independently of chromosome instability. We identify that cells with centrosome amplification are primed for apoptosis. We show they are dependent on the apoptotic inhibitor BCL-XL, and that this is not a consequence of mitotic stresses associated with centrosome amplification. Given the multiple mechanisms that promote chemotherapy responses in cells with centrosome amplification, we assess such a relationship in an epithelial ovarian cancer patient cohort. We show that high centrosome numbers associate with improved chemotherapy responses and longer overall survival. Our work identifies apoptotic priming as a clinically relevant consequence of centrosome amplification, expanding our understanding of this pleiotropic cancer cell feature.

## INTRODUCTION

Centrosomes are the major microtubule organizing centers in proliferating animal cells, whose structure and number are tightly regulated during the cell cycle (*1*). The centrosome is duplicated during S-phase in a PLK4 kinase dependent manner and the two centrosomes contribute to the timely and functional assembly of a bipolar spindle during mitosis. In cancer cell lines, centrosome structural and numerical defects are common (*2*) and in particular centrosome amplification – more than 2 centrosomes per cell - has also been observed *in situ* in tumor samples (*3–5*). Cells with centrosome amplification perform bipolar mitosis via centrosome clustering mechanisms during spindle assembly (*6–10*). Centrosome amplification is nevertheless associated with chromosome instability (*11, 12*), and increased invasive behaviors (*13–15*). As first postulated by T. Boveri (*16*), centrosome amplification can drive tumorigenesis *in vivo* (*7, 17–19*). Defects in centrosome clustering capacity are associated with lethal multipolar divisions (*8, 11, 20*), motivating the search for inhibitors that limit centrosome clustering and eliminate cells with centrosome amplification (*20–25*).

The combination of Carboplatin and Paclitaxel is used as standard of care in various cancers including ovarian, breast, and lung cancer. Carboplatin induces DNA damage and Paclitaxel stabilizes microtubules leading to cell death via mitotic catastrophe which is defined as death during or following abnormal mitosis (*26, 27*). Despite the central role of centrosomes in spindle assembly, how centrosome amplification influences the response to combined Carboplatin and Paclitaxel remains unexplored. Paclitaxel has been shown to induce multipolar divisions (*28*) and this can be favored by centrosome amplification (*29*). The impact of centrosome amplification on the response to DNA damaging agents has however not been explored despite centrosomes regulating multiple signaling pathways that could influence chemotherapy responses (*30–34*). Multiple consequences of centrosome amplification could therefore synergize with combined Carboplatin and Paclitaxel to induce efficient cancer cell elimination.

Here we chose to study how centrosome amplification influences the response to combined Carboplatin and Paclitaxel in the context of epithelial ovarian cancer, a disease with poor clinical outcome related to late diagnosis, and frequent relapse (*35*). Centrosome amplification is observed in ovarian cancer cell lines, and we recently also characterized its occurrence *in situ* in patient samples (*5*). We use an inducible PLK4 over-expression system in ovarian cancer cell lines to induce centrosome amplification in isogenic backgrounds. We perform single-cell live-imaging of cells to assess the correlations between mitotic behaviors and cell fate during chemotherapy. We show that centrosome amplification favors the response to combined Carboplatin and Paclitaxel via multiple mechanisms. Beyond multipolar divisions associated with Paclitaxel exposure, we found that centrosome amplification also favors cell death independently of mitotic behaviors. We show that centrosome amplification, although well tolerated by ovarian cancer cells, leads to mitochondria outer membrane permeabilization priming. We assess the level of centrosome amplification in a previously characterized ovarian cancer patient cohort and observe an association between high centrosome numbers and the patient time to relapse as well as their overall survival. Together our work shows for the first time that centrosome amplification can synergize with combined chemotherapy, completing our understanding of its consequences in cancer.

## RESULTS

### Centrosome amplification favors cell death in response to combined Paclitaxel and Carboplatin

To study the influence of centrosome amplification on the response to Carboplatin and Paclitaxel, we used an inducible PLK4 over-expression system (PLK4OE) in the epithelial ovarian cancer cell line OVCAR8. Exposing cells to doxycycline for 72h at 1µg/mL induced centrosome amplification (more than 2 centrosomes per cell) in around 80% of cells, compared to 4% in control OVCAR8 (DMSO treated, PLK4Ctl) (Fig. S1A-B). We used MTT viability assays to determine Carboplatin and Paclitaxel IC50s over the 72h following PLK4OE induction (Fig. S1C-D). For both drugs, the IC50 was lower for PLK4OE (67μM and 3,4nM respectively) compared to PLK4Ctl (136μM and 5,1nM respectively), already suggesting that centrosome amplification sensitizes cells to these chemotherapeutic agents. We also determined working combination concentrations (100μM Carboplatin + 3,3nM Paclitaxel) that induce 60% growth inhibition in PLK4Ctl and 89% in PLK4OE (Fig. S1E).

We next used live-imaging to investigate how centrosome amplification influences the response to chemotherapy and in particular if it favors catastrophic mitosis. We used H2B-RFP expressing OVCAR8 allowing us to observe chromosome behaviors during mitosis and chromatin compaction that occurs when cells die (Fig. 1A). We imaged cells during the 72h of exposure to 100μM Carboplatin + 3,3nM Paclitaxel and performed analysis of complete cell lineages, counting the number of cells produced per lineage (proliferation) and the fate of these cells (viability). In untreated OVCAR8 cells, PLK4OE reduced proliferation compared to PLK4Ctl, but independently of an increase in cell death (Fig. 1B). Combined chemotherapy reduced proliferation of both PLK4Ctl and PLK4OE cells and this reduction was associated with an increase in cell death (Fig. 1B) as well as cell cycle lengthening (Fig. S1F). In agreement with the MTT dose-response assays, combined chemotherapy induced a stronger reduction of viable cells produced per lineage in PLK4OE compared to PLK4Ctl, and this was associated with a higher proportion of cell death with 75% dying in PLK4OE and 33% dying in PLK4Ctl (Fig. 1B and Fig. 1E-F). By examining cell fate in consecutive generations, we observed that cell death was mainly occurring in generations 2 and 3 suggesting that passage through mitosis or extended exposure time to chemotherapy is detrimental for the progeny (Fig. 1C).

**Fig. 1.**
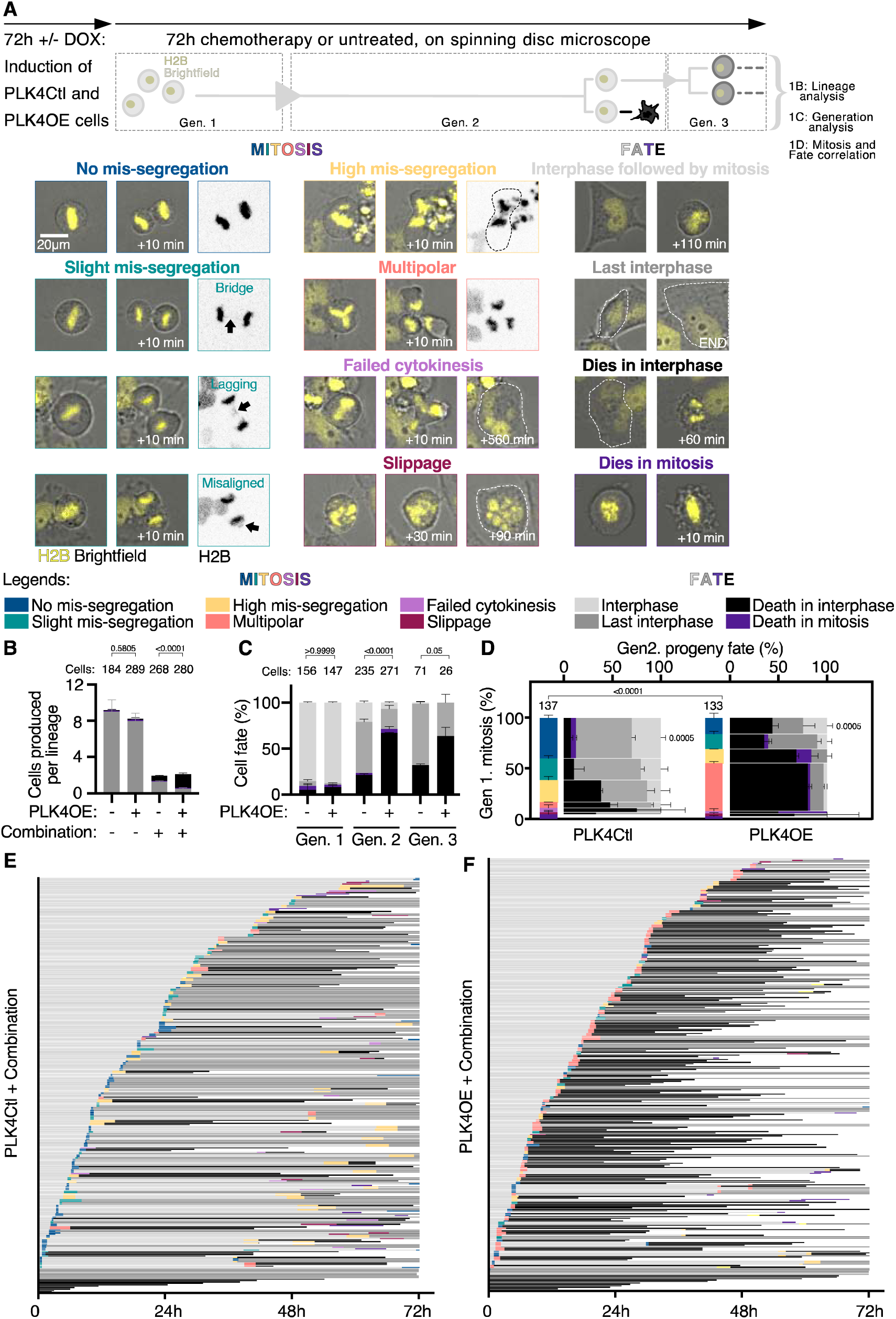
Centrosome amplification favors cell death in response to combined chemotherapy. **(A)** Single-cell live-imaging workflow. OVCAR8 cells expressing H2B-RFP and inducible for PLK4 over-expression are exposed to DMSO or 1µg/ml Doxycycline (DOX) for 72h to induce centrosome amplification. PLK4Ctl and PLK4OE cells are then filmed during 72h of chemotherapy and lineages are tracked over multiple generations. Representative images of mitotic behaviors and cell fates are shown with the color-coded legends used in the subsequent panels. Lineage analysis consists in counting for each starting cell, the number of cells adopting the different fates (See panel B). Generation analysis consists in determining the percentage of a generation that will adopt the different fates (See panel C). Mitosis and fate correlation consists in determining the percentage of cells adopting the different fates, depending on the behavior of the mother cell during mitosis (See panel D). **(B)** Bar graphs showing the averages and SEM of the number of cells per lineage adopting the indicated fates (Legends in panel A). A minimum of 20 lineages were analyzed from two independent experiments, statistical tests: Fisher’s exact test on the number of cell death events (pooling death in interphase and in mitosis). Numbers on the top of each graph represent the number of cells analyzed per condition. **(C)** Bar graphs showing the average and SEM of the percentages of cells undergoing indicated fates (Legends in panel A). Two independent experiments, statistical tests: Fisher’s exact test on the number of cell death events (pooling death in interphase and death in mitosis). Numbers on the top of each graph represent the number of cells analyzed per condition. **(D)** Vertical axis: Bar graphs showing the average and SEM of the percentages of mitotic phenotypes (Legends in panel A). 137 and 133 cell divisions were analyzed from two independent experiments, statistical test: Fisher’s exact test on the number of multipolar divisions. Horizontal axis: Bar graphs showing the average and SEM of the percentages of cells undergoing indicated fates (Legends in panel A) according to the mitotic behavior of mother cells, with bar width depending on the proportion of cells displaying a given mitotic phenotype. Two independent experiments, statistical test: Fisher’s exact test on the number of *No mis-segregation* progeny (progeny of blue mitosis) dying in mitosis and interphase. **(E- F)** Single cell profiles of PLK4Ctl (E) and PLK4OE (F) undergoing Carboplatin+ Paclitaxel exposure. Each row corresponds to one cell (Legends in panel A).

We therefore characterized the mitotic behaviors in the first generation focusing on chromosome mis-segregation (Fig. 1A). In untreated PLK4Ctl OVCAR8, we observed a significant proportion of chromosome instability with around 20% of mitosis occurring with either chromosome mis-alignment, one lagging chromosome, or one chromatin bridge (Fig. 1A black arrows and S1G and I). These behaviors, which are sometimes difficult to discriminate at the low spatio-temporal resolution used in our long-term live-imaging approach, were pooled together and considered as *Slight mis-segregation* events (Fig. 1A). These events were more frequent in PLK4OE (36%), in agreement with centrosome amplification inducing merotelic attachments (*11*). However, the proportion of *Multipolar* divisions was negligible, suggesting these cells are competent to cluster supernumerary centrosomes (Fig. S1G-I). Importantly, combined chemotherapy induced an increase in two behaviors associated with strong chromosome mis-segregation: *Multipolar* divisions and bipolar divisions associated with multiple chromosome mis-segregation events (combinations of bridges, lagging and misaligned chromosomes – termed here *High mis-segregation*) (Fig. 1A and D vertical axis, Fig. S1G-H). Events of complete division failure – either via cytokinesis failure, mitotic slippage, or death in mitosis – were observed but remained infrequent upon combined chemotherapy (Fig. 1D vertical axis). PLK4OE cells exposed to combined chemotherapy present close to 47% of multipolar divisions compared to 6% observed in PLK4Ctl (Fig. 1D vertical axis and Fig. 1E-F). We next focused on the fate of the progeny produced by the different cell division categories and observed that within cells completing cell division, higher levels of chromosome mis-segregation were associated with increased cell death in the progeny (Fig. 1D horizontal axis). In particular, multipolar divisions were associated with at least 50% cell death in the progeny, and the increase of these multipolar divisions in PLK4OE cells exposed to combined chemotherapy therefore contributes to the decreased viability observed in this condition. Paclitaxel has been suggested to induce multipolar spindles in cells that present centrosome amplification (*29*), and exposing PLK4OE cells to Paclitaxel alone also induces multipolar divisions (Fig. S1J), suggesting the increased multipolarity we observe in presence of the combined chemotherapy is caused by the effect of Paclitaxel on the capacity of cells to cluster centrosomes. However, and surprisingly, we also observed that independently of the type of mitosis induced by combined chemotherapy, the proportion of cell death in the progeny was higher in PLK4OE compared to PLK4Ctl (Fig.1D, horizontal axis). This was in particular the case for the progeny of cells that do not show any chromosome mis-segregation, where 40% cell death is observed in PLK4OE compared to only 12% cell death in PLK4Ctl (Fig. 1D, horizontal axis). These results suggest that centrosome amplification favors cell death in response to combined chemotherapy independently of multipolarity and chromosome segregation errors.

### Centrosome amplification favors cell death in response to Carboplatin independently of catastrophic mitosis

We were next interested in understanding why cell death is enhanced in PLK4OE in response to combined chemotherapy irrespective of mitotic behaviors. We observed that the IC50 of Carboplatin is lower for PLK4OE compared to PLK4Ctl (Fig. S1D). We therefore hypothesized that PLK4OE cells may respond differently to DNA damage induced by Carboplatin. In both cell populations, Carboplatin exposure at the IC50 determined for PLK4Ctl (136μM) induced an increase in DNA double-strand breaks, as visualized via staining for γ-H2AX, an early marker of the DNA damage response (Fig. S2H-I). γ-H2AX fluorescence intensity was similar in Plk4OE and PLK4Ctl, suggesting centrosome amplification does not increase the levels of DNA damage in response to Carboplatin. We therefore performed live-imaging to better understand how OVCAR8 cells respond to 136μM Carboplatin, and how centrosome amplification modifies this response. First focusing on the lineage analysis, we observed that Carboplatin treated cells have a reduction in proliferation compared to untreated cells, and an increase in cell death (Fig. S1F, Fig. 2A-D) . In agreement with PLK4OE cells being more sensitive to Carboplatin, fewer viable cells were produced in Carboplatin exposed PLK4OE compared to PLK4Ctl, and this was associated with an increase in cell death with 47% in Plk4OE and 30% in PLK4Ctl (Fig. 2A-D). Similar to the combined chemotherapy, most cell death occurred in generation 2, suggesting that mitosis contributes to cell death in response to Carboplatin (Fig. 2E). We therefore focused on chromosome mis-segregation during the first mitosis. In both PLK4Ctl and PLK4OE treated with Carboplatin, the main phenotype was an increase in *High mis-segregation* divisions, and unlike in the response to combined chemotherapy, *Multipolar* divisions remained negligible (Fig. 2A-B and F). The observed increase in cell death in PLK4OE is therefore completely independent of the capacity to assemble a bipolar spindle.

**Fig. 2.**
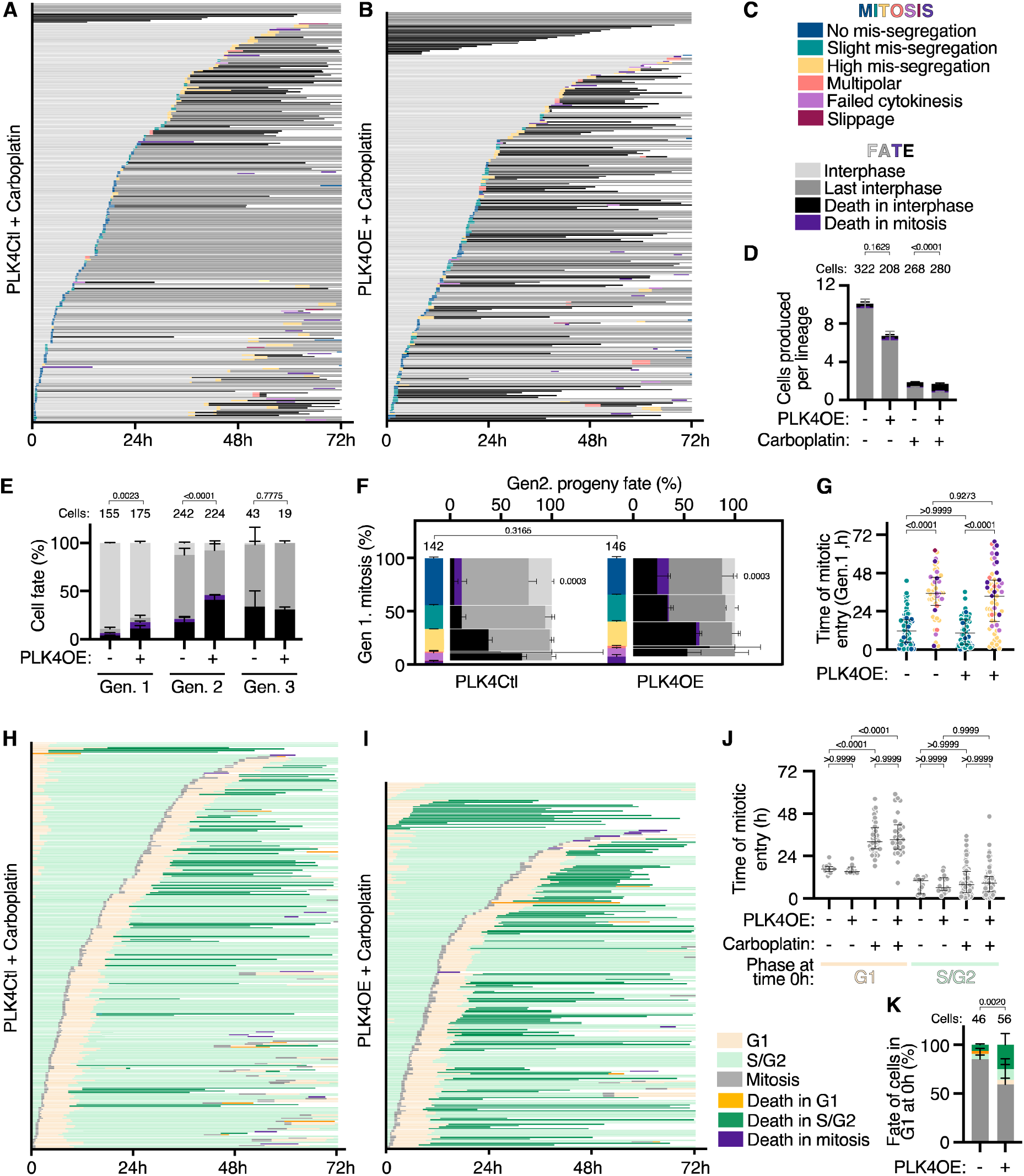
Centrosome amplification enhances cell death in response to Carboplatin independently of chromosome mis-segregation. **(A- B)** Single cell profiles of PLK4Ctl (A) and PLK4OE (B) undergoing Carboplatin exposure. Color coding of mitosis and fates legends in panel C. **(C)** Legends for panels A,B,D,E,F and G, as defined in Figure 1A. **(D)** Bar graphs showing the averages and SEM of the number of cells produced per lineage, adopting the indicated fates (Legends in panel 1A). A minimum of 31 lineages were analyzed from two independent experiments, statistical tests: Fisher’s exact test on the number of cell death events (pooling death in interphase and in mitosis). **(E)** Bar graphs showing the average and SEM of the percentages of cells undergoing indicated fates (Legends in Figure 1A and panel C). Two independent experiments, statistical tests: Fisher’s exact test on the number of cell death events (pooling death in interphase and death in mitosis). Numbers on the top of each graph represent the number of cells analyzed per condition. **(F)** Vertical axis: Bar graphs showing the average and SEM of the percentages of mitotic phenotypes (Legends in Figure 1A and panel C). 142 and 146 cell divisions analyzed from two independent experiments, statistical test: Chi-square test. Horizontal axis: Bar graphs showing the averages and SEM of the percentages of cells undergoing indicated fates (Legends in Figure 1A and panel C) according to the mitotic behavior of the mother cell, with bar width depending on the proportion of cells according to their the mitotic behavior. Two independent experiments, statistical test: Fisher’s exact test on the number of *No mis-segregation* progeny (progeny of blue mitosis) dying in mitosis and interphase. **(G)** Scatter dot plot graphs showing time of mitotic entry, with median and interquartile range. Cells were classified depending on mitotic phenotypes with color-code defined in panel C. Two independent experiments with a minimum of 48 mitosis analyzed per category. Statistical tests: Kruskal-Wallis with Dunn’s multiple comparisons tests. **(H- I)** Single cell profiles of FUCCI PLK4Ctl (I) and PLK4OE (I) cells undergoing Carboplatin exposure. Times in G1, S/G2 and mitosis, as well as death in each of these phases are color-coded as indicated. **(J)** Scatter dot plots of the time of mitotic entry depending on cell-cycle phase at movie start, with median and interquartile range. Two independent experiments with a minimum of 10 times analyzed per category. Statistical tests: Kruskal-Wallis with Dunn’s multiple comparisons tests. **(K)** Bar graphs showing the average and SEM of the percentage of cells adopting the indicated fates (legends in panel I). Two independent experiments, statistical test: Fischer’s exact test on the number of cells dying (irrespective of the cell-cycle phase). Numbers on the top of each graph represent the number of cells analyzed per condition.

Sorting single cell behaviors based on the time of first mitotic entry reveals that in PLK4Ctl, *high mis-segregation* events occur during mitosis initiated after a longer time spent in Carboplatin (Fig. 2A and G). It is then mainly the progeny from these divisions that died (Fig. 2A and F). To better understand this time-dependent-response, we analyzed the cell-cycle in response to Carboplatin using a FUCCI expressing OVCAR8 cell-line. This strategy allowed us to discriminate cells in G1 from cells in S/G2 (Fig. 2H-I and Fig. S2A-B), while still observing cell death events which occur mainly in S/G2 (Fig. S2E). First, we observed that Carboplatin induced an increase in S/G2 length, suggesting DNA damage in OVCAR8 cells activates the intra S and/or the G2/M checkpoint (Fig. S2C), while G1 length was not strongly varying in any observed condition (Fig. S2D). Next, we observed that S/G2 lengthening and subsequent delayed mitotic entry occurred mainly in cells that were in G1 at the onset of Carboplatin exposure, and therefore in cells exposed to Carboplatin for a complete S-phase (Fig. 2H and J). Despite this delay, these cells most likely entered mitosis with unrepaired damage, driving high levels of chromosome mis-segregation during mitosis and eventually leading to death of the progeny. In PLK4OE cells exposed to Carboplatin, the timing and proportions of mitotic and cell-cycle behaviors was similar to PLK4Ctl. *High mis-segregation* events occured in cells with a strong delay in mitotic entry (Fig. 2B, F and G), which concurred with cells that were in G1 at the onset of Carboplatin exposure (Fig. 2I and J). However, in PLK4OE, the association between cell death and mitotic phenotypes was different from PLK4Ctl. Indeed around 35% of cell death was observed for the progeny of cells that had *No mis-segregation* or *Slight mis-segregation* defects, in contrast with PLK4Ctl where less than 10% of these cells died (Fig. 2F, horizontal axis).

These observations therefore suggest that PLK4Ctl cells are essentially killed by catastrophic mitosis induced by high levels of DNA damage, while PLK4OE cells do not require a catastrophic mitosis to be eliminated. In agreement with cell death occurring independently of mitosis in PLK4OE, 17% cells died in the first generation compared to only 6% in PLK4Ctl (Fig. 2A-B and E). In particular, cells that were in G1 at the beginning of Carboplatin exposure and therefore can accumulate DNA damage in their first cell-cycle, were preferentially killed in PLK4OE with 25% of cells dying compared to 8% in PLK4Ctl (Fig. 2H-I and K). Our findings propose that centrosome amplification sensitizes cells to the effect of carboplatin in a single cell-cycle and independently of catastrophic mitotic behaviors.

The centrosome has previously been involved in regulating the DNA damage response via recruitment of ATR, ATM, Chk1 and Chk2 (*30, 31*). We investigated whether centrosome amplification modifies the signaling downstream of DNA damage, explaining increased cell-death in response to Carboplatin. We detected phosphorylation and activation of Chk1 and p53 in response to Carboplatin, but no difference is observed in PLK4OE compared to PLK4Ctl cell extracts (Fig. S2F-G). In agreement with DNA damage levels and responses being unchanged by PLK4OE, we also observed no difference in the recruitment of DNA damage repair factors FANCD2, 53BP1 and Rad51 (Fig. S2H-I). Together, these observations suggest that centrosome amplification favors cell death in response to DNA damage, independently of catastrophic mitosis, but also independently of the DNA damage response.

### Centrosome amplification modulates mitochondrial apoptosis independently of p53 and the PIDDosome

In order to better understand how centrosome amplification favors cell death in response to Carboplatin independently of mitotic errors, we characterized the type of cell death and the associated signaling network. We observed that Carboplatin treatment induced apoptosis characterized by Caspase-3 cleavage (Fig. 3A-B), as well as cells becoming positive for Annexin-V by flow cytometry, which was completely suppressed by the pan-caspase inhibitor Q-VD-Oph (Fig. 3C). PLK4OE induces both a premature cleavage of Caspase-3 (t=48h compared to t=72h in PLK4Ctl), as well as an increase in the Annexin-V positive cell population (53% compared to 32% in PLK4Ctl), confirming that centrosome amplification favors the apoptotic response to Carboplatin (Fig. 3A-C). To determine if the mitochondria outer membrane permeabilization (MOMP) dependent apoptotic pathway was activated in response to Carboplatin, we stained cells for Cytochrome C in order to observe its release from mitochondria (Fig. 3D). In untreated cells, Cytochrome C was detected in punctate structures throughout the cytoplasm, in agreement with its mitochondrial localization. Upon Carboplatin exposure, the Cytochrome C staining remained similar to untreated cells, while we observed dead cell remnants characterized by condensed DNA (Fig. 3D White arrows and 3E). However if release of Cytochrome C occurred, it could lead to immediate apoptosis initiation and detachment of the cells, precluding the observation of cells via immunofluorescence. We therefore use Q-VD-Oph to inhibit apoptosis in response to Carboplatin and observed a population of cells where Cytochrome C is diffuse in the cytoplasm and nucleus (Fig. 3D Pink arrows and 3E). MOMP, Cytochrome C release and caspase activation therefore occurrs in response to Carboplatin, suggesting that mitochondrial apoptosis is the main cell death mechanism at play.

**Fig 3.**
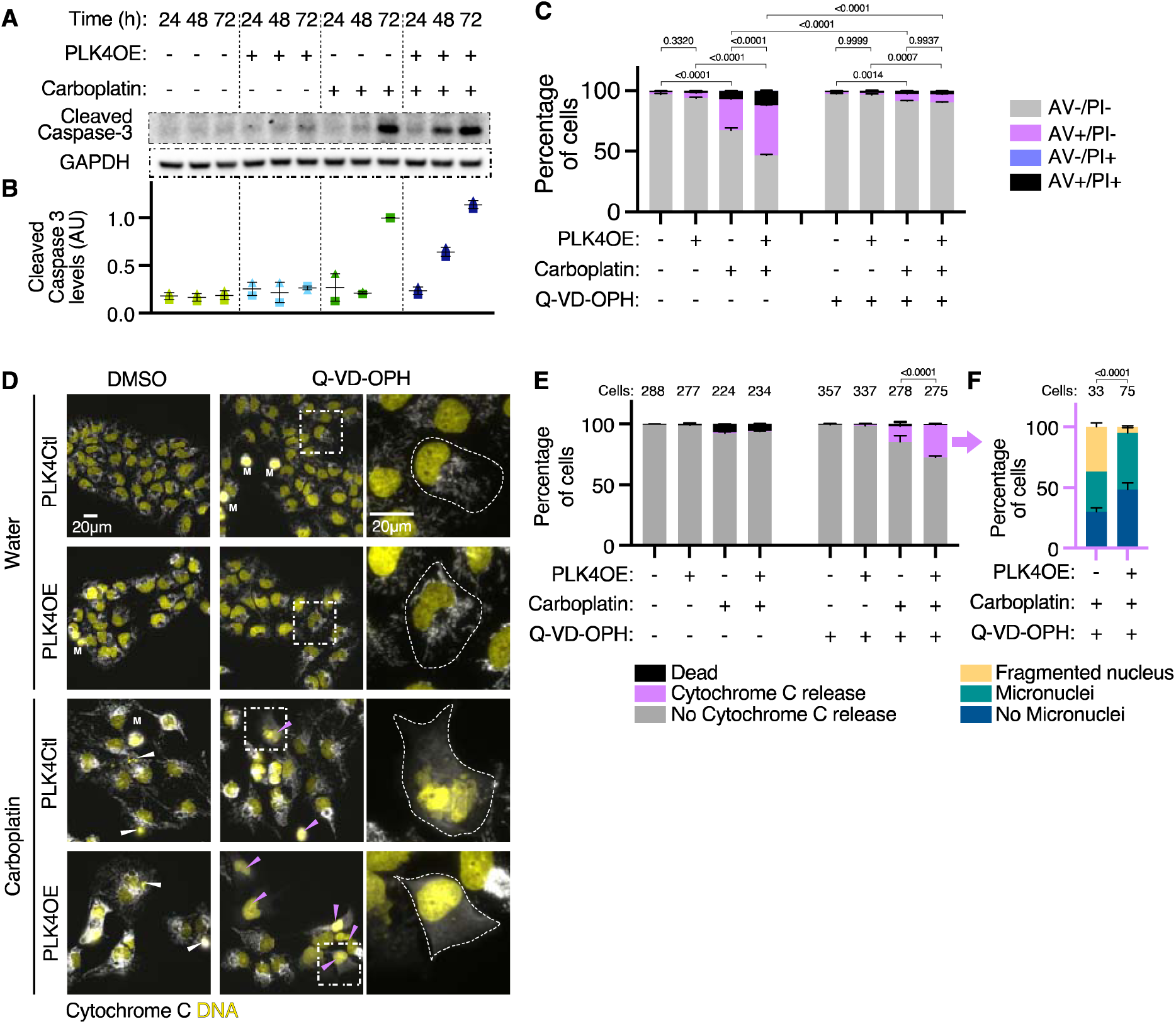
Centrosome amplification favors mitochondrial outer membrane permeabilization in response to Carboplatin. **(A)** Representative Western Blot detecting Caspase-3 cleavage. **(B)** Average and SEM of cleaved caspase 3 protein levels from 2 independent experiments, normalized to levels measured in Carboplatin treated PLK4Ctl cells at 72h. **(C)** Bar graphs showing the average and SEM of the percentage of cells in specified Annexin V-APC/PI gates analyzed by flow cytometry. 4 replicates obtained from 2 independent experiments, with a minimum of 60000 cells analyzed per condition and replicate. Statistical test: comparison of the percentage of Annexin V positive cells, using ANOVA with Sidak’s multiple comparison test. Representative cytometry profiles can be found in the supplementary materials. **(D)** Representative images of cells labelled with DAPI (yellow) and antibodies against Cytochrome C (gray). White arrows indicate dead cell debris, pink arrows indicate cells that have released Cytochrome C in the cytoplasm, M indicates mitotic cells. Representative insets are shown for individual Q-VD-Oph treated cells. **(E)** Bar graphs showing the average and SEM of the percentages of indicated cell populations. Two independent experiments, statistical test: Fisher’s exact test on the number of cells releasing Cytochrome C. Numbers on the top of each graph represent the number of cells analyzed per condition. **(F)** Bar graphs showing the average and SEM of the percentages of cells releasing Cytochrome C with indicated nuclear phenotypes. Two independent experiments, statistical test: Fisher’s exact test on the number of cells with fragmented nuclei. Numbers on the top of each graph represent the number of cells analyzed per condition.

Importantly, we observed 27% of cells releasing Cytochrome C in PLK4OE cells compared to only 12% in PLK4Ctl cells, suggesting that Carboplatin induces a stronger activation of apoptosis in the presence of centrosome amplification (Fig. 3D-E). In agreement with different stresses leading to apoptosis between PLK4Ctl and PLK4OE, we observed that 36% of cells poised to die with fragmented nuclei in PLK4Ctl compared to 5% in PLK4OE (Fig. 3F). These fragmented nuclei are symptomatic of high chromosome mis-segregation during mitosis, supported by PLK4Ctl cells being killed by catastrophic mitosis, while centrosome amplification favors apoptosis independently of catastrophic mitosis in PLK4OE.

The canonical intrinsic apoptosis pathway linking the DNA damage response to MOMP occurs via p53 stabilization which then drives the transcription of pro-apoptotic BCL2 family genes (*36*). Centrosome amplification has been linked to p53 stabilization via PIDDosome activation, which is dependent on centriole distal appendage grouping. This leads to Caspase-2 cleavage and activation and cleavage of MDM2 - a major p53 regulator – (*33, 34*). OVCAR8 cells have a mutant TP53 gene which leads to alternative splicing of exon5, and a 6 amino-acid deletion in p53’s DNA binding domain (*37*). We observed that p53 protein is present, phosphorylated, and accumulates in response to Carboplatin (Fig. S2F, quantified in Fig. S3B). Although the deletion in the DNA damage binding domain is suggested to preclude its transcriptional activities (*37*), we observed that its transcriptional targets p21 and PUMA mildly increase upon Carboplatin exposure (Fig. S3A-B). Using shRNA, we knocked-down TP53 and showed that p53 was dispensable for cell death in both PLK4Ctl and PLK4OE in response to Carboplatin (Fig. S3C-D). p21 is best characterized for its functions in cell-cycle arrest and apoptosis inhibition, but has also been shown to upregulate apoptosis (*38*). As we noticed p21 to be upregulated in PLK4OE cells compared to PLK4Ctl (Fig. S3A-B), we tested its contribution to cell death, but observed no effect of knocking down the p21 coding gene CDKN1a (Fig. S3C- D). Despite apoptosis being p53 independent in Carboplatin treated OVCAR8, we tested whether PIDDosome activation may contribute to enhancing intrinsic apoptosis in PLK4OE. Indeed, upon PLK4OE, we observed cleavage of Caspase-2 and MDM2 reflecting PIDDosome activation (Fig. S3E-F). We knocked-down the distal appendage protein required for PIDDosome activation ANKRD26 and observed a strong reduction of Caspase-2 and MDM2 cleavage in PLK4OE, reflecting efficient PIDDosome silencing (Fig. S3G). However, this had no effect on the enhanced cell death observed upon Carboplatin exposure in PLK4OE, suggesting that centrosome amplification favors apoptosis independently of the PIDDosome (Fig. S3H). Altogether, our results show that centrosome amplification leads to enhanced apoptosis in response to Carboplatin independent of previously described centrosome signaling nodes.

### Centrosome amplification primes for MOMP and sensitizes cells to a diversity of chemotherapies

So far we have observed that PLK4OE cells execute apoptosis faster and to a higher proportion, in response to a level of stress (mitotic behaviors, DNA damage levels and DNA damage response) which is no different than in PLK4Ctl. This therefore suggested that these cells may be primed for MOMP, meaning that the balance between pro-apoptotic and anti-apoptotic BCL2 family proteins that determine the activity of the mitochondrial pore forming proteins BAX and BAK is tilted towards their activation in PLK4OE (*39*). To test this possibility, we performed MTT dose-response assays during 72h to drugs which mimic the activity of specific pro-apoptotic BH3-only BCL2 family proteins. Strikingly, we observed that PLK4OE induces a strong sensitization to WEHI-539 - a specific BCL-XL inhibitor - (Fig. 4A-B), but not to Venetoclax or A1210477 - specific inhibitors of BCL-2 and MCL-1 respectively- (Fig. S4A-B). We confirmed that 72h 300nM WEHI-539 exposure selectively induced apoptosis in PLK4OE cells in a Caspase dependent manner, via Annexin V and PI cytometry (Fig. 4C). We also observed release of Cytochrome C from mitochondria in WEHI-539 treated PLK4OE cells upon pan-caspase inhibition, confirming that BCL-XL inhibition induces MOMP specifically in PLK4OE (Fig. 4D-E). Counting centrosomes in PLK4OE cells revealed that the 72h 300nM WEHI-539 treatment reduced the proportion of cells with extra centrosomes to the level observed in PLK4Ctl cells (Fig. 4F-G). These results suggest efficient killing of cells with centrosome amplification, which is suppressed upon pan-caspase inhibition. We were also able to count centrosomes in the population of cells that release Cytochrome C from mitochondria and are therefore poised to die, revealing that the majority of these cells have centrosome amplification in WEHI-539 treated PLK4OE cells (Fig. 4G). Interestingly, an extremely small proportion (1%) of PLK4Ctl cells also release Cytochrome C in response to WEHI-539 (Fig. 4E) and counting centrosomes in these cells revealed a higher level of centrosome amplification than the untreated PLK4Ctl population (Fig. 4G). This suggests that independently of induced PLK4 over- expression, centrosome amplification primes for MOMP in OVCAR8 cells. We confirmed this observation in parental OVCAR8 cells devoid of the inducible PLK4OE transgene to exclude the possibility that PLK4OE leakage sensitizes the PLK4Ctl cells to WEHI-539 inhibition (Fig. S4C).

**Fig 4.**
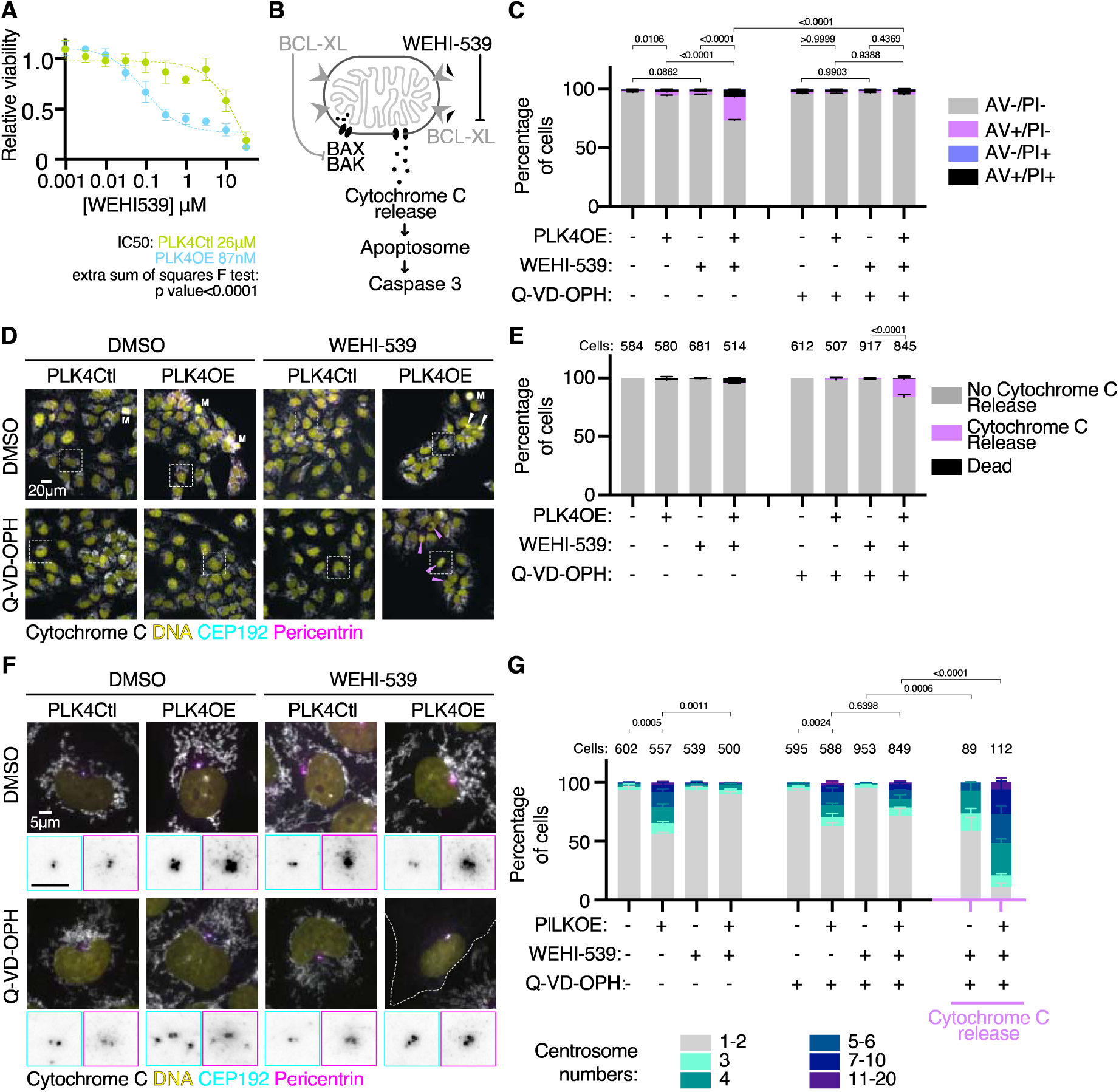
Centrosome amplification primes for mitochondria outer membrane permeabilization. **(A)** Dose-response of PLK4Ctl and PLK4OE cells to WEHI-539, normalized to their respective untreated conditions, obtained from MTT viability assays. Mean and SEM of 3 independent experiments each obtained from averaging 3 technical replicates. **(B)** Schematic of the induction of apoptosis by WEHI-539. When BAX and BAK channel formation are inhibited by BCL-XL, Cytochrome C is present in the mitochondria intermembrane space (Left). WEHI-539 inhibits BCL-XL which relieves the inhibition of channel formation by BAX and BAK, leading to Cytochrome C release, Apoptosome activation, and cleavage of Caspase 3 (Right). **(C)** Bar graphs showing the average and SEM of the percentage of cells in specified Annexin V-APC/PI gates analyzed by flow cytometry. 4 replicates obtained from 2 independent experiments with a minimum of 15000 cells analyzed per condition and replicate. Statistical test: comparison of the percentage of Annexin V positive cells, using ANOVA with Sidak’s multiple comparison test. Representative cytometry profiles can be found in the supplementary materials. **(D)** Representative images of cells labeled with DAPI (yellow) and antibodies against Cytochrome C (gray), CEP192 (Cyan) and Pericentrin (Magenta). White arrows indicate dead cell debris, pink arrows indicate cells that have released Cytochrome C in to the cytoplasm, M indicates mitotic cells. Representative insets are shown in panel (F). **(E)** Bar graphs showing the average and SEM of the percentages of indicated cell populations. Two independent experiments, statistical test: Fisher’s exact test on the number of cells releasing Cytochrome C. Numbers on the top of each graph represent the number of cells analyzed per condition. **(F)** Insets from panel (D) with inverted grayscale insets zooming on the centrosomes showing CEP192 (cyan border) and Pericentrin (Magenta border). **(G)** Bar graphs showing the average and SEM of the percentage of cells with the indicated number of centrosomes (determined by the co-localization of CEP192 and Pericentrin). Two independent experiments, statistical test: comparison of the percentage of cells with more than 2 centrosomes, using ANOVA with Sidak’s multiple comparison test. Numbers on the top of each graph represent the number of cells analyzed per condition.

We next tested if similar effects might be observed in other cell lines and established inducible centrosome amplification via PLK4OE in ovarian cancer cell lines COV504 and SKOV3 (Fig. S4D-E). We then tested if MOMP priming identified in OVCAR8 was also observed in these cell lines (Fig. S4F). We used the less specific BH3-mimetic Navitoclax (inhibitor of BCL2, BCL-XL and BCL2), as the dependency on BCL-XL in PLK4OE OVCAR8 might be reflecting OVCAR8 apoptotic wiring rather than a specific effect of centrosome amplification on BCL-XL. We observed that Navitoclax reduced the viability of PLK4OE cells preferentially compared to PLK4Ctl cells in COV504 (EC80 of 400nM and 3,6uM respectively) although to a lesser extent than in OVCAR8 (EC80 of 40nM and 2,2uM respectively). Priming was however not observed in SKOV3. We then established the IC50s for Carboplatin and Paclitaxel, and determined combination concentrations in PLK4Ctl cells (Fig. S4G). We used Trypan blue assays to determine viability in response to chemotherapy and confirmed this approach in OVCAR8 by showing that viability decreases more in PLK4OE compared to PLK4Ctl (Fig. S4H left). Interestingly, we observed a gradation in the enhanced cell death induced by centrosome amplification in the different cell lines, with the strongest effect observed in OVCAR8 (Fig. S4H left), the weakest in SKOV3 (Fig. S4H right), and an intermediate effect in COV504 (Fig. S4H middle), which can be put in perspective with the observed gradation in MOMP priming. For responses to Paclitaxel however we have to take into account the fact that multipolar divisions are also favored in COV504 PLK4OE cells (Fig. S4I).

Our identification of apoptotic priming in cells with centrosome amplification suggests that it might be associated with enhanced cell death in response to a larger panel of drugs. We therefore tested whether centrosome amplification sensitizes ovarian cancer cells to PARP inhibitors which are now included in standard of care protocols in epithelial ovarian cancer (*40*), focusing on Olaparib (IC50s determined and presented in Fig. S4G). Trypan Blue viability assays in OVCAR8 and COV504 confirmed that PLK4OE leads to reduced viability compared to PLK4Ctl in response to Olaparib (Fig. S4J). This effect was not observed in SKOV3 in which we have not observed MOMP priming linked to centrosome amplification. Together our results suggest that centrosome amplification enhances cell death in response to chemotherapy differentially depending on the cell line, and that centrosome amplification associated apoptotic priming can sensitize to a diversity of chemotherapies.

### Centrosome amplification primes for MOMP independently of chromosome instability or lengthened mitosis

Centrosome amplification induces an increase in chromosome instability and a spindle-assembly checkpoint dependent extension of mitosis duration (*7, 8, 11*), both of which we observed in PLK4OE OVCAR8 cells (Fig. S1F and S1I). Apoptotic priming and in particular sensitization to BCL-XL inhibition has previously been linked to mitotic defects. In particular chromosome instability, micronuclei formation and cGAS/STING signaling can drive a transcriptional response that drives apoptosis or priming (*41, 42*). Alternatively, extended mitotic duration can lead to the proteosomal degradation of anti-apoptotic BCL2 family proteins, leading to BCL-XL sensitization (*43–46*). We were therefore interested in determining if apoptotic priming observed in response to centrosome amplification is caused by cumulated mitotic stress in these cells.

First, we aimed to better characterize the mitotic stress induced by centrosome amplification in the already chromosomally instable OVCAR8 cell line (Fig. S1I). We used single-cell DNA-sequencing to assess karyotype heterogeneity (Supplementary Material and Methods) and observed scores of 0,119 in PLK4Ctl, 0,137 in PLK4OE cells and 0,283 in PLK4Ctl cells treated with 1μM of the MPS1 inhibitor AZ3146 as a positive control of chromosome mis-segregation (Fig. 5A). PLK4OE therefore only mildly increased aneuploidy, in line with levels of chromosome-mis-segregation observed by time-lapse imaging (Fig. 5B). We also assessed the extent of mitotic lengthening induced in PLK4OE cells (Fig. 5C) and observed it was mild (median=60min in PLK4OE and median=35min in PLK4Ctl) compared to that induced by low doses of the CENP-E inhibitor GSK923295 (median=100min at 30nM and 175min at 35nM), despite levels of chromosome mis-segregation being similar (Fig. 5B).

**Fig. 5.**
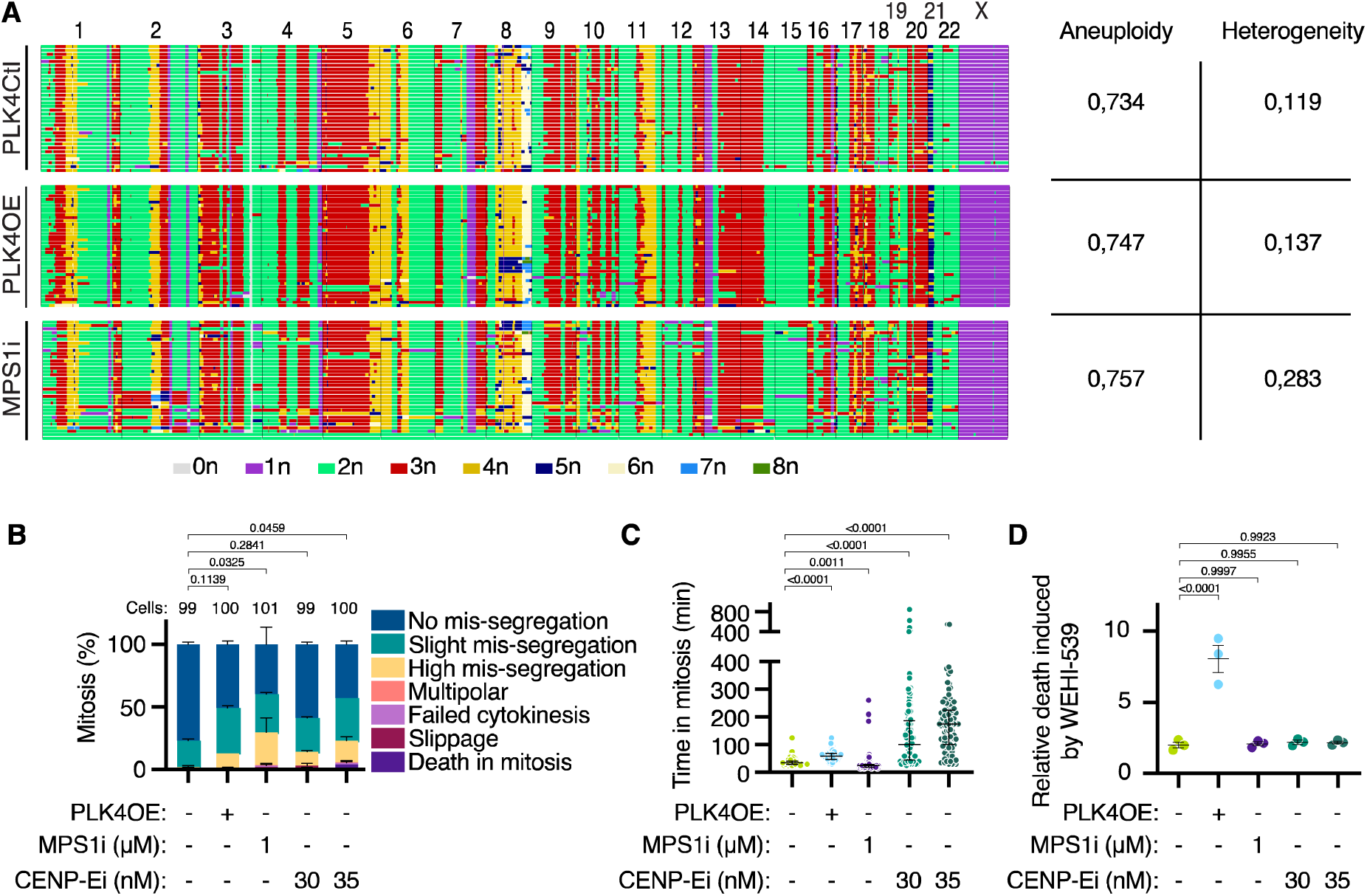
Centrosome amplification primes for mitochondria outer membrane permeabilization independently of mitotic stress. **(A)** Genome-wide copy-number plots for G1 OVCAR8 cells. Each row represents a cell. Indicated aneuploidy and heterogeneity scores are calculated as described in material and methods. **(B)** Bar graphs showing the average and SEM of the percentage of mitotic phenotypes as defined in Figure 1A. Two independent experiments, statistical test: comparison of the percentage of cells with no mis-segregation using ANOVA with Dunnett’s multiple comparison test. Numbers on the top of each graph represent the number of cells analyzed per condition. **(C)** Scatter dot plot graph of mitosis length with median and interquartile range. At least 96 mitosis analyzed from two independent experiments, statistical test: Kruskall-Wallis with Dunn’s multiple comparison test. **(D)** Bar graphs showing the ratio between the percentages of Annexin V positive cells observed in presence and absence of WEHI-539 300nM. Average and SEM of 3 replicates from two independent experiments. Statistical test: ANOVA with Dunnett’s multiple comparison test. Numbers on the top of each graph represent the number of cells analyzed per condition.

Mitotic stress is therefore mild in PLK4OE compared to the other perturbations we tested, but we were nevertheless interested in determining if it contributes to apoptotic priming. We were not able to reduce mitotic duration in PLK4OE cells via spindle assembly checkpoint inhibition without inducing a strong increase in multipolar divisions (data not shown), so we used MPS1 and CENP-E inhibition to mimic mitotic stress observed in PLK4OE. We pretreated PLK4Ctl cells with inhibitors during 72h before adding WEHI-539 for an additional 24h. In PLK4OE cells this lead to 33% Annexin V positive cells, whereas it only induces 9% in response to MPS1 inhibition (Fig. S5A), making it unlikely that priming occurs in response to chromosome instability in PLK4OE. In CENP-E inhibition pretreated cells at 30nM and 35nM, WEHI-539 induces 12% and 25% Annexin V cells respectively, in line with mitotic lengthening inducing priming (*46*). Importantly however, 35nM CENP-E inhibition pretreatment already induces 12% Annexin V positive cells which is considerable compared to the 4% observed in PLK4OE (Fig. S5A), and most likely is explained by the extensive mitotic lengthening observed in response to 35nM CENP-E inhibition (Fig. 5C). This results in the ratio of cell death induced by WEHI-539 relative to the basal observed level of cell death to be comparable in PLK4Ctl cells and CENP-E inhibition pretreated cells (around 2-fold). In contrast, the ratio of cell death was much higher and close to 8 fold increase upon PLK4OE (Fig. 5D). Therefore, the priming induced in PLK4OE stands out from that induced by other sources of mitotic stress in that PLK4OE cells are viable but strongly dependent on BCL-XL. These results also suggest that the combination of chromosome instability and mitotic lengthening is not the major contributor to MOMP priming upon centrosome amplification in OVCAR8 cells.

To identify transcriptomic signatures that may influence cell death responses in cells with extra centrosomes, we used bulk RNAseq comparing PLK4OE and PLK4Ctl OVCAR8 cells. A strong inflammatory signature in PLK4OE (Fig. S5B) was identified, and we also observed STING phosphorylation (Fig. S5C), suggesting that the cGAS/STING pathway may shape the transcriptional response to centrosome amplification. We were therefore interested in directly testing if cGAS/STING signaling might contribute to priming, although this seemed unlikely as CENP-E and MPS1 inhibition also activate STING (Fig. S5C) but are not associated with priming. We used a bulk LentiCRISPR knock-out approach of STING, but observed no influence on PLK4OE cells sensitivity to WEHI-539 (Fig. S5D-E).

We therefore identify that centrosome amplification in OVCAR8 leads to MOMP priming which is revealed by a selective sensitization to the BCL-XL inhibitor WEHI-539. Comparing with other mitotic perturbations, we conclude that the centrosome amplification associated priming is independent of mitotic lengthening and chromosome instability.

### High centrosome numbers are associated with a better response to chemotherapy in a High grade Serous Ovarian Cancer patient cohort

To assess if centrosome amplification is associated with chemotherapy responses in patients, we turned to the characterization of centrosomes we previously performed *in situ* in treatment-naive epithelial ovarian tumors. Here centrosomes were detected as the colocalization of Pericentrin and CDK5RAP2 in confocal images of methanol-fixed patient tissue sections (*5*). To assess centrosome numbers in samples we defined the centrosome to nucleus ratio (CNR) as the number of centrosomes detected in a field by the number of nuclei which we averaged over 10 fields per patient (Fig. 6A). In healthy tissues obtained from prophylactic oophorectomy or hysterectomy, the CNR was 1.02±0.02 suggesting cells have on average one centrosome per cell which is expected for a non-proliferative tissue. On average, the CNR in tumor tissues was 1.43±0.04, with the minimum at 0.61 and maximum at 2.55. While only 9% of tumors had a CNR above 2, suggesting that pervasive centrosome amplification - when defined by the presence of more than 2 centrosomes per cell - is infrequent, 89% of the tumors presented a CNR superior to the average CNR found in healthy tissues. Considering that the CNR did not correlate with proliferation as established by the mitotic index (Fig. S6A) and Ki67 staining (Fig. S6B), centrosome amplification could contribute to this increase in CNR in tumors compared to healthy samples.

**Fig. 6.**
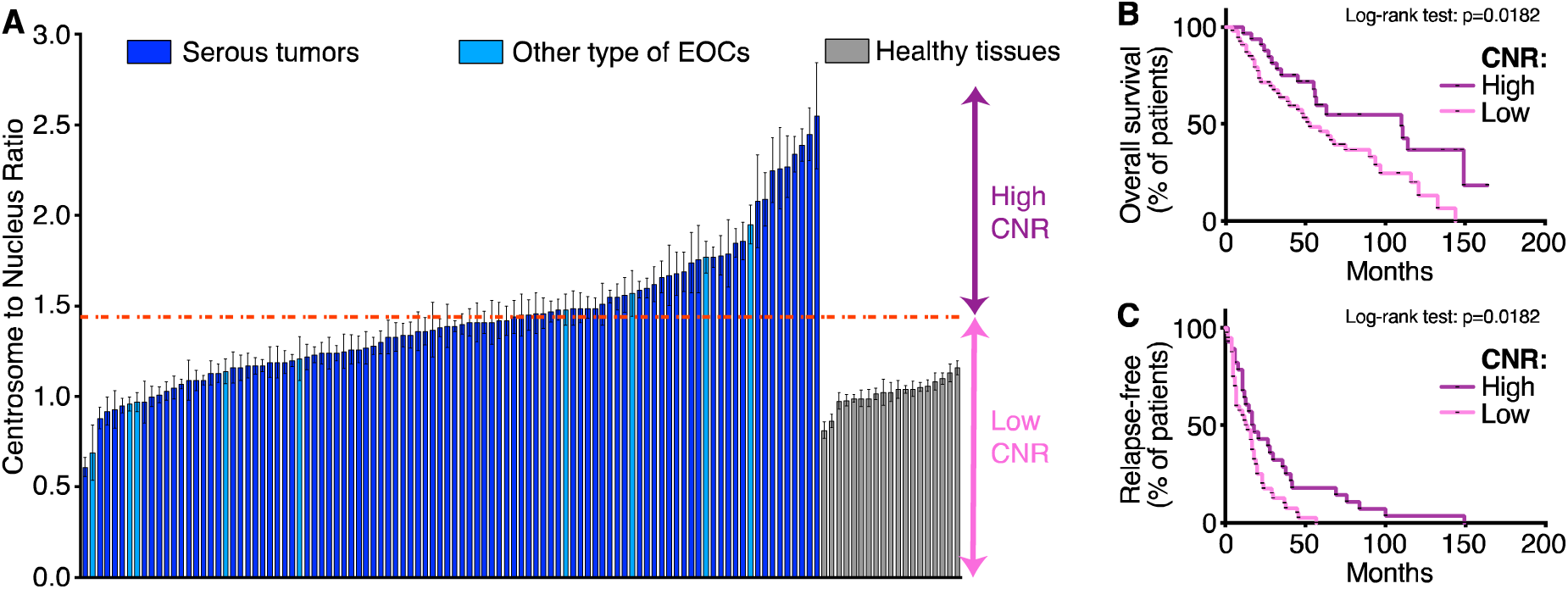
High centrosome numbers favor the response to chemotherapy in a HGSOC patient cohort. **(A**) Average and SEM of the CNR established in 10 fields per tumor or healthy tissue sample. Red dotted line indicates CNR=1,45, the cut-off between High-CNR and Low-CNR patients. **(B-C)** Kaplan-Meier curves for overall survival (B) and relapse-free time after the first line of chemotherapy (C) according to CNR status.

We next examined if the CNR was associated with chemotherapy responses restricting our analysis to the high-grade serous ovarian cancers (HGSOCs) in our cohort. We dichotomized our population into two groups using the Classification And Regression Trees (CART) method, resulting in the categorization of the cohort into 55 low CNR (<=1.45) tumors and 33 high CNR (> 1.45) tumors. Importantly, we observed no association between CNR and FIGO stage, with 59% stage III patients in the cohort comprising both High and Low CNR (Fig. S6C). We next plotted HGSOC patient survival curves according to the CNR status. We found that high CNR was associated with better overall survival (Fig. 6B). These results suggest that despite its oncogenic potential (*7, 17–19*), centrosome amplification might improve patient prognosis at least in ovarian cancer. This puzzling observation could be explained if High CNR promotes chemotherapy responses as overall survival data reflects patients complete clinical course which includes Carboplatin and Paclitaxel therapy in this cohort. To directly assess a link between CNR and chemotherapy responses, we plotted patient time to relapse and found that High CNR was associated with a longer time to relapse (Fig. 6C). Together, this work suggests that centrosome status in ovarian cancer can influence patient outcome, in particular with High CNR favoring response to chemotherapy.

## DISCUSSION

Centrosome amplification as a therapeutic target has been mainly explored from the prism of multipolar division, but mitotic drugs that limit centrosome clustering have had limited success in the clinic (*48*). Our results identify apoptotic priming as a novel cell death susceptibility conferred by centrosome amplification. In particular, we show that centrosome amplification sensitizes cells to BH3-mimetic drugs.

The apoptotic priming seems to be specific to centrosome amplification, rather than a consequence of the associated mitotic stress. Possible causes of this priming could be disruption of mitochondrial networks during mitosis, or in interphase. This may be in link with recent observations of subcellular reorganization in response to centrosome amplification in RPE-1 cells (*49*), although we observe no striking effect on mitochondria organization in OVCAR8 (data not shown). Centrosomes are also involved in multiple signaling pathways (*31, 50*) and given the pleiotropic effects of centrosome amplification which also favors ROS and inflammation (*15*), we consider that the best method to identify the source of the priming would be whole-genome screening approaches.

From a clinical perspective, our analysis of a patient cohort shows that high centrosome numbers limit relapse in response to chemotherapy, indicating that centrosome amplification must be considered beyond its malignant potential. Given the toxicity of cytotoxic therapies, the perspective of better patient stratification and response prediction, considering centrosome amplification as a sensitizing factor offers promising perspectives. Our observation that centrosome amplification enhances cell death independently of multipolar mitosis broadens the therapeutic importance of this cancer cell feature, beyond treatments that target spindle assembly and mitosis Our identification of apoptotic priming in cells with centrosome amplification also has clinical relevance, and asks whether centrosome numbers could be considered to better direct BH3-mimetic administration whose use in the clinic is hampered by lack of good prognostic markers (*51*). Alternatively, we also wonder if centrosome amplification might be a contributor to the on-target toxicity of BCL-XL inhibition leading to thrombocytopenia. Indeed, the megakaryocytes that produce platelets are polyploid and present multiple centrosomes (*52*) and this might contribute to their targeting by BCL-XL inhibitors.

There are multiple limitations to our study. We must emphasize that the levels of centrosome amplification in the cohort is low (*5*) and that centrosome loss might also contribute to modulating chemotherapy responses. It nevertheless remains interesting to consider that targeting low levels of centrosome amplification could have an observable clinical effect, and to explain these results we propose that elimination of cells with centrosome amplification might be advantageous given the malignant potential of these cells (*7, 13–15, 19*). We are also eager to know if centrosome numbers influence responses to chemotherapy in additional epithelial ovarian cancer cohorts and in different cancer types. An important step to facilitate broader studies is the automatization of centrosome detection and counting in patient tissues. Additionally our identification of apoptotic priming in response to centrosome amplification and the associated clinical perspectives justify the need for a better understanding of the priming mechanism induced by centrosome amplification. This would also help identify the contexts in which this priming emerges as we have not observed it in all the cancer cell lines studied.

## MATERIAL AND METHODS

### Study design

This work is a study of the influence of centrosome amplification on the response to chemotherapy in epithelial ovarian cancer. The objectives of the cell biology work were to identify if and how centrosome amplification favors cell death in response to chemotherapy, using a combination of single cell live imaging, and classical cell biology experiments such as cytometry and Western Blot. All the presented data has been replicated in 2 to 5 biological replicates. The objective of the clinical work was to identify if centrosome numbers influence clinical parameters in a patient cohort. All samples were taken before chemotherapy administration and obtained from the Biological Resource Center (BRC) of Institut Curie (certification number: 2009/33837.4; AFNOR NF S 96 900). In compliance with the French regulation, patients were informed of the studies performed on tissue specimens and did not express opposition. The National Commission for Data Processing and Liberties (N° approval: 1487390) approved all analysis, as well as The Institutional Review Board and Ethics committee of the Institut Curie. Centrosomes were previously stained and detected in a HGSOC patient cohort of 100 patients (*5*). Here we determined an index allowing the quantitative assessment of centrosome numbers in patient tissues, which we then correlated with patient clinical parameters. Data collection for each experiment is detailed in the respective figure legend.

### Cell lines and cell culture

All cell lines were cultured at 37°C with 5% CO2 in DMEM/F12 media (ThermoFisher Scientific #31331028) supplemented with 10% Tetracyclin-free Fetal Bovine Serum (Dutscher #500101L), 100 µg/ml streptomycin and 100U/ml penicillin (ThermoFisher Scientific #15140122). OVCAR8 and COV504 were obtained from the laboratory of F. Mechta-Grigoriou, and SKOV3 were purchased from ATCC (#HTB-77). Cell cultures underwent authentification by short tandem repeat analysis (powerplex16 HS kit, Promega #DC2101) and were routinely checked for mycoplasma (PlasmoTest Mycoplasma detection kit, InvivoGen, #rep-pt1).

### Cell line generation

Inducible PLK4 over-expression, H2B-RFP expression, FUCCI expression, shRNA expression and bulk CRISPR-Cas9 knock-out of STING were stably established by lentiviral infection. Viruses were produced in HEK cells using Lipofectamine 2000 (ThermoFisher Scientific #11668019) to co-transfect lentiviral constructs with pMD2.G and psPAX2 plasmids. Viral particles were collected in the supernatant 48h after transfection, filtered, and used to infect the cell lines during 24h. Cells were then FACS sorted (inducible PLK4 over-expression and H2B- RFP) or selected using Puromycin at 5 µg/ml (shRNA lines and CRISPR-Cas9 knock-out of STING). Efficiency of knock-down and knock-out was assessed by Western Blot. The list of plasmids used is available in Table S1.

### Drug treatments

All chemicals used are listed in Table S2. To induce centrosome amplification, cells were exposed to doxycycline (1 µg/ml) or DMSO (diluent control, 1/10000) for 72h. If cells were subsequently treated with another drug, cells were detached and replated without addition of doxycycline to the PLK4OE population, and left to attach for 8h. Drug treatments were then carried out for 72h at the indicated concentrations. For the experiments comparing centrosome amplification to other mitotic stresses (CENP-E and MPS1 inhibition), cells were exposed to doxycycline (1 µg/ml for centrosome amplification), AZ3146 (1µM, for MPS1 inhibition), GSK923295 (30-35nM, for CENP-E inhibition) or DMSO (diluent control) for 72h. Subsequent treatments (WEHI-539) or analysis (live-imaging of mitotic phenotypes) were then carried out in presence of the same initial concentrations of drug for 24h.

### Live-Imaging and analysis

For live-imaging of chemotherapy responses, cells were plated on Ibidi µ-Slide 8 Well slides (Clinisciences, #80806-G500). Chemotherapy treated and untreated cells from both PLK4Ctl and PLK4OE populations were imaged during the same experiment. Imaging was performed with a 20x objective (CFI Plan Apo LBDA 20x 0,75/1 mm CCo 0,17) via an EMCCD camera (Evolve, Photometrics) on an inverted microscope (Inverted Ti-E Nikon) equipped with a spinning disk (CSU-X1 Yokogawa), a stage-top temperature and CO2 incubator (Tokai Hit) and integrated in Metamorph software. For each well, 4-6 positions were acquired every 10min during 72h, with a single slice in the brightfield channel and 10 slices per z-stack in the H2B-RFP channel or in the mKO2-Cdt1(30–120) and mAzami-Green-Gem1(1–110) channels for the FUCCI cells. Time-lapse movies were then analyzed manually using a custom Fiji macro to record a list of events, and a custom Python script to generate excel data files and single-cell profiles.

For live-imaging of mitotic phenotypes induced by centrosome amplification, MPS1 inhibition and CENP-E inhibition, the same approach was used, acquiring each position every 5min during 24h. Imaging was performed using the equipment described above. .

### Immunofluorescence

Cells were plated on 18mm glass coverslips in 12-well plates. Cells were fixed for 5min in ice-cold methanol (for Fig.s S1A and S4D), or for 10min in 4%PFA at room-temperature (for Fig.s 3D, 4C, 4E) or at 4°C (for Fig. S2H). Cells were washed 3 times in PBST (PBS + 0,1% Triton X- 100) and incubated in PBST + BSA 0,5% for 30min at room temperature. Cells were then incubated for 1h in primary antibodies diluted in PBST + BSA 0,5%, washed 3 times in PBST, incubated for 30min in secondary antibodies diluted in PBST + BSA 0,5%, and washed 3 times in PBST. Cells were then stained for DNA using 3 µg/ml DAPI diluted in PBST + BSA 0,5%, washed 3 times in PBS and mounted with mounting medium (1.25% n-propyl gallate, 75% glycerol, in H2O). Antibodies used are listed in Table S3.

### Immunofluorescence imaging and quantifications

Immunofluorescence images were acquired with a sCMOS camera (Flash 4.0 V2, Hamamatsu) on a widefield microscope (DM6B, Leica systems), with a 63x objective (63x HCX PL APO 1.40-0.60 Oil from Leica), using Metamorph software. Z-stacks were acquired at 0,3 µm intervals.

Centrosome numbers and cytochrome C release were scored manually. DNA damage marker intensity or foci number were determined on z-projections of images, using a custom Python script to run the h_maxima function from the skimage.morphology.extrema module.

### Western Blotting

Cells were lysed in RIPA (150mM sodium chloride, 1% NP-40, 0.5% sodium deoxycholate, 0.1% sodium dodecyl sulfate, 50mM Tris, pH8.0) complemented with protease (Sigma-Aldrich # 11697498001) and phosphatase (Sigma-Aldrich #4906845001) inhibitors. Samples were dosed using a BiCinchoninic acid Assay (Pierce BCA protein assay, ThermoFisher Scientific # 23227). Samples were diluted in RIPA with 4X NuPage LDS sampling buffer (ThermoFisher Scientific # NP0007) and heated at 80°C for 10min. 20µg of protein was loaded in Bolt 4-12% Bis-Tris precast gels (ThermoFisher Scientific #NW04125BOX), and subjected to electrophoresis in Bolt MOPS SDS running buffer (ThermoFisher Scientific # B0001). The gels were transferred to nitrocellulose membranes (Dutscher # 10600001) using transfer buffer (25mM Tris, 192mM Glycine, 20% Methanol) for 90min at 4°C. Membranes were stained in primary or horse-radish peroxidase coupled secondary antibodies diluted in PBS or TBS + 0,5% Tween 20 + 0,5% BSA or non-fat milk according to providers instructions. Membranes were first stained using Ponceau, before saturating for 1h at room temperature in 5% non-fat dry milk or 5% BSA in PBS or TBS + 0,5% Tween20. Membranes were then incubated overnight in primary antibodies, washed 5 times in PBS or TBS + 0,5% Tween 20, then incubated for 1h at room temperature in secondary antibodies. Membranes were then washed again 5 times in PBS or TBS +0,5% Tween 20. Horse-radish Peroxidase reaction was developed using SuperSignal Plus Chemiluminescent substrates (Thermo Fisher Scientific # 34580 and #34094) and imaged (BioRad ChemiDoc MP). The Image Lab software (BioRad version 6.0.1) was used to measure background-adjusted volume intensity, which was normalized using GAPDH signal. Antibodies used are listed in Table S3.

### Transfection

HT-DNA was transfected as a positive control for cGAS/Sting activation. Trasnfection of 1µg/mL HTDNA was carried out using Lipofectamine 2000 (ThermoFisher Scientific #11668019) for 24h.

### Cytometry

Cells were detached, rinsed in PBS, rinsed in AnnexinV Binding Buffer (BioLegend # 422201), and around 100000 cells were resuspended in 50µL Annexin V Binding Buffer. Cells were stained using Annexin V APC and Propidium Iodide (Biolegend # 640932) at 0,6μg/mL and 10mg/mL respectively. Cells were incubated for 15min, then diluted in 200μL Annexin V Binding Buffer. Cells were analysed using a Bio-Rad ZE5 analyzer, and data was analyzed using FlowJo 10.6.0 software.

### MTT viability assays

For dose-response to drugs, cell viability was assessed using MTT viability assays. Cells were plated in triplicates at 15000cells/well in 96-well plates and left for 2h to adhere prior to drug addition. Cells were left to grow for 72h, and MTT diluted in PBS was added at 5μg/mL. After 4h incubation, medium was removed and replaced by 150uL DMSO. 570nm absorbance was performed on a BMG Labtech ClarioStar plate reader. Triplicates were averaged and normalized by untreated controls.

### Trypan Blue Proliferation and Viability assays

For proliferation and viability assays, cells were plates at 100000 cells/well in 6-well plates. Cells were then detached, resuspended in 500uL medium, and live/dead cells were counted using a Beckman Coulter Vi-Cell cell counter.

### RNA sequencing

Following centrosome amplification with doxycycline for PLKOE vs DMSO for PLK4Ctl, total RNA was extracted with RNeasy Mini kit (Qiagen #74104) following manufacturer’s instructions. RNA integrity and quality were checked with Agilent RNA 6000 Nano Kit (Agilent, # 5067-1511) and corresponding devices. Samples were processed at Institut Curie NGS platform from cDNA synthesis, amplification, quality assessment and sequencing. Novaseq 6000 system (Illumina) was used for sequencing (read length of 100 bp, paired end) . All the bioinformatic analysis were done by Genosplice (http://www.genosplice.com) including quality control of sequences generated, read mapping and gene differential analysis (R software, Deseq2). Biological interpretation of the identified genes was done using GSEA tool for pathway enrichment analysis between distinct conditions.

### Single-cell whole genome sequencing

Cells were treated with DMSO (1/10000), Doxycycline (1µg/mL) or AZ3146 (1µM) for 72h. Cells were then frozen in freezing medium (10% DMSO, 40% FBS in DMEM-F12).

#### Nuclei preparation and sorting

Cells were thawed, and single-cell sequencing was performed on cell nuclei isolated from cell lysis, leaving the nucleus intact. Thawed cells were prepared by resuspending in PBS + 1% BSA, washing, and pelleting. To generate nuclei, cells were resuspended and incubated (15 minutes on ice in dark environment) in cell lysis buffer (100 mM Tris-HCl pH 7.4, 154 mM NaCl, 1 mM CaCl2, 500 µM MgCl2, 0.2% BSA, 0.1% NP-40, 10 µg/mL Hoechst 33358, 2 µg/mL propidium iodide in ultra-pure water). Resulting cell nuclei were gated for G1 phase (as determined by Hoechst and propidium iodide staining) and sorted into wells of 96 wells plates on a MoFlo Astrios cell sorter (Beckman Coulter), depositing one cell per well. 96 wells plates containing nuclei and freezing buffer were stored at −80°C until further processing. Automated library preparation was then performed as previously described (*53*).

#### AneuFinder analysis

Sequencing was performed using a NextSeq 2000 machine (Illumina; up to 120 cycles; single end or up to 113 and 7 cycles; paired end). The generated data were subsequently demultiplexed using sample-specific barcodes and changed into fastq files using bcl2fastq (Illumina; version 1.8.4). Reads were afterwards aligned to the human reference genome (GRCh38/hg38) using Bowtie2 (version 2.2.4; (*54*)). Duplicate reads were marked with BamUtil (version 1.0.3; (*55*)).

The aligned read data (bam files) were analyzed with a copy number calling algorithm called AneuFinder (version 1.14.0; (*56*)) using an euploid reference (*57*). Following GC correction (R package: BSgenome.Hsapiens.UCSC.hg38_1.4.1; The Bioconductor Dev Team 2015) and blacklisting of artefact-prone regions (extreme low or high coverage in control samples), libraries were analyzed using the dnacopy and edivisive copy number calling algorithms with variable width bins (average binsize = 1 Mb; step size = 500 kb) and breakpoint refinement (refine.breakpoints = TRUE). Results were afterwards curated by requiring a minimum concordance of 90 % between the results of the two algorithms. Libraries with on average less than 10 reads per bin and per chromosome copy (∼ 55,000 reads for a diploid genome) were discarded.

#### Aneuploidy score

The aneuploidy score of each bin was calculated as the absolute difference between the observed copy number and the expected copy number when euploid. The score for each library was calculated as the weighted average of all the bins (size of the bin as weight) and the sample scores were calculated as the average of the scores of all libraries.

#### Heterogeneity score

The heterogeneity score of each bin was calculated as the proportion of pairwise comparisons (cell 1 vs. cell 2, cell 1 vs cell 3, etc.) that showed a difference in copy number (e.g. cell 1: 2- somy and cell 2: 3-somy). The heterogeneity score of each sample was calculated as the weighted average of all the bin scores (size of the bin as weight).

### Centrosome numbers in tumors

For each sample, 10 randomly chosen fields were considered. Using ImageJ software, we visually counted the number of nuclei and the number of centrosomes in a blind manner without taking into account tumor identity. The Centrosome to Nuclei Ratio (CNR) was obtained by dividing the total number of centrosomes by the total number of nuclei in each field.

### Proliferation and mitotic index

For Ki67 proliferation assessment, we performed immunochemistry assays using mouse anti-human ki67 antibody (M7240, DAKO, 1/200 at pH9) in a series of paraffin-embedded tissue blocks of HGSOC. Sections of 3 µm were cut using a microtome from the paraffin-embedded tissue blocks of normal tissue and invasive lesions. Tissue sections were deparaffinized and rehydrated through a series of xylene and ethanol washes. Briefly, the key steps included: (i) antigen retrieval with ER2 pH9, (Leica: AR9640); (ii) blocking of endogenous peroxidase activity with Bond polymer refine detection kit (Leica: DS9800) (iii) incubation with primary antibodies against the targeted antigen; (iv) immunodetection with Revelation and counter staining Bond polymer refine detection kit (Leica: DS9800). Immunostaining was performed using a Leica Bond RX automated immunostaining device. We performed an immunohistochemical score (frequency x intensity) through analysis of 10 high-power fields (HPF, x 400). All quantifications were performed by 2 pathologists with blinding of patient status.

For mitotic index, paraffin-embedded tissue sections of tumors were stained with hematoxylin and eosin. The mitotic count was determined by the number of mitotic figures found in 10 consecutive high-power fields (HPF), in the most mitotically active part of the tumor (entire section). Only identifiable mitotic figures were counted. Hyperchromatic, karyorrhectic, or apoptotic nuclei were excluded.

### Statistical analysis

Statistical analysis were performed using GraphPad Prism. The tests used are specified in the figure legends. The numbers of cells analyzed and the number of replicates are reported either on the figure or in the respective figure legends.

## List of Supplementary Materials

Figs. S1 to S6

Table S1 to S3

Representative cytometry profiles

## Acknowledgments

We are grateful to the patients who consented to participate in this research and to the medical teams involved in their care. We thank Andrew Holland for the kind gift of the inducible PLK4 over-expression plasmid. We thank the ICGEX platform at Institut Curie (IC) headed by Sylvain Baulade for bulk RNA sequencing and Genosplice (http://www.genosplice.com) and P. Delagrange and W. Zeitouni for performing the analysis. We thank the Tissue Imaging (PICT- IBiSA) and Nikon Imaging Centre at IC, member of the French National Research Infrastructure France-BioImaging (ANR10-INBS-04) and the IC cytometry platform. We thank the Genomics Platform of the translational research department at IC for cell line authentication. We thank Nicolas Manel and Sebastian Montealegre for discussions and the gift of reagents used to characterize cGAS/STING signaling. We thank Stephen Taylor for discussions about this project. We also thank all the members of the Basto team for discussions and comments on the manuscript.

## Author contributions

The project was designed and conceptualized by FE, and RB with significant input from OG and GF. FE, GF, and AYS performed most experiments including time-lapse imaging, image analysis, cytometry, immunofluorescence, western blots and quantifications. FE developed the code used for time-lapse imaging of single-cell responses to chemotherapy. GF prepared cells for single cell whole genome sequencing which were then processed by AET, with data analysis performed by RW under the supervision of DCJS and FF. AYS and OG prepared cells for bulk RNA sequencing, and OG supervised the data analysis. SG helped establish the stable FUCCI cell lines; AH contributed to determine IC50s; SF helped analyze time-lapse imaging experiments. XS-G and AVS provided human samples from the pathology department of IC. OM managed the patient database, and SRR provided advice in methodology. JPM quantified the centrosomes in patient tissue samples and OG supervised their analysis. Figures were prepared by FE with help from GF and AYS. FE wrote the manuscript with input from RB and OG. The work was supervised by FE, OG and RB.

## Data and materials availability

All data and code are available upon request.

## SUPPLEMENTARY MATERIAL

### Supplementary Figures

**Fig. S1.**
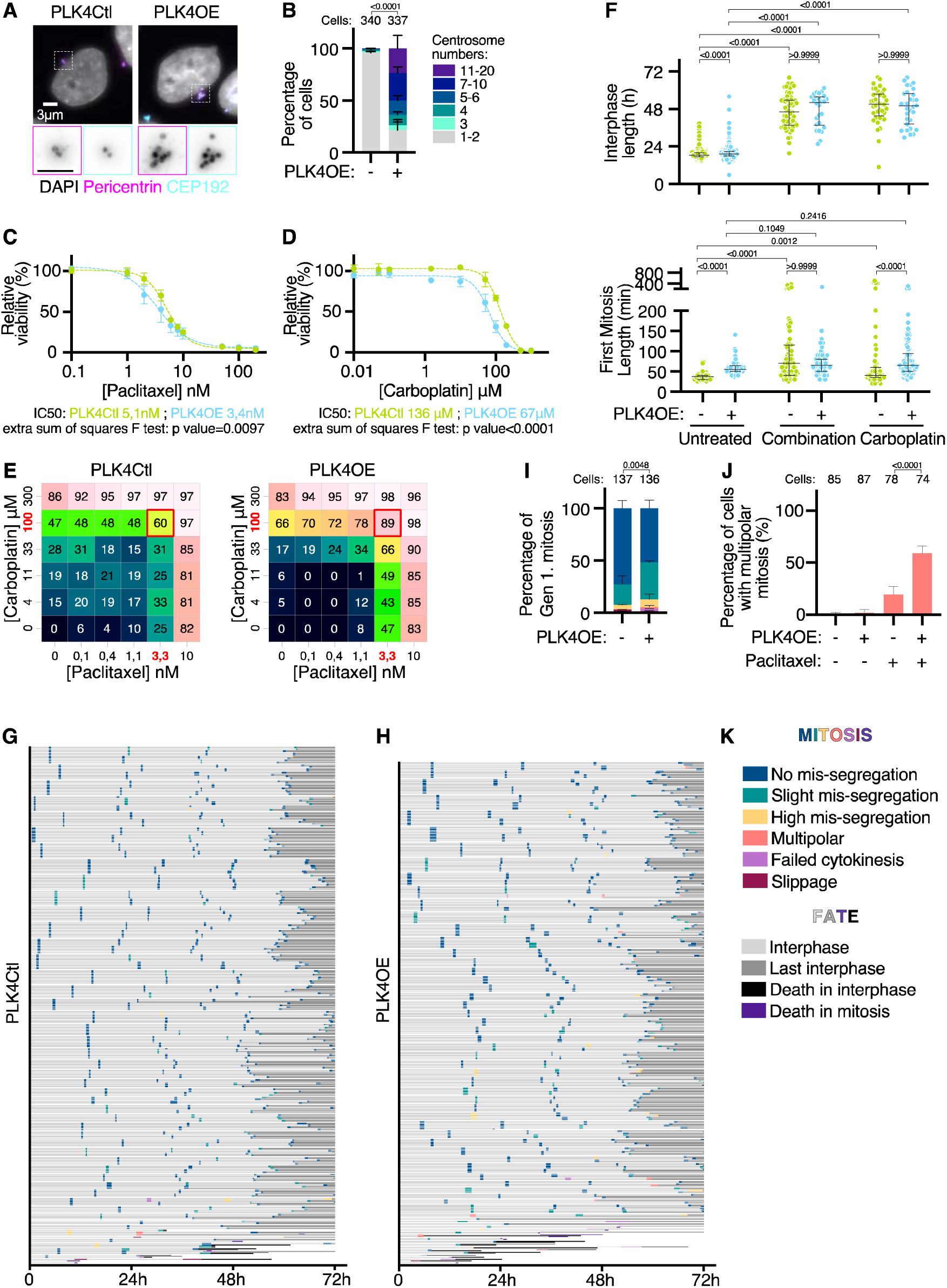
**(A)** Representative images of OVCAR8 cells stained with DAPI (gray) and antibodies against CEP192 (Cyan) and Pericentrin (Magenta). **(B)** Bar graphs showing the averages and SEM of the percentage of cells with the indicated number of centrosomes (CEP192 dots colocalizing with Pericentrin). 3 independent experiments, statistical test: Fisher’s exact test comparing the number of cells with more than 2 centrosomes. **(C- D)** Dose-response of PLK4Ctl and PLK4OE cells to Paclitaxel (C) and Carboplatin (D), normalized to their respective control conditions, obtained from MTT viability assays. Mean and SEM of 2 independent experiments each obtained from averaging 3 technical replicates. **(E)** Combination matrixes for Carboplatin and Paclitaxel combined treatment, representing percentage of viability inhibition compared to control cells. Chosen working concentrations are highlighted in red. **(F)** Scatter dot plots of Interphase length (top) and First mitosis length (bottom), with Median and interquartile range. Data from two independent experiments are pooled for Combined treatment and Carboplatin treatment, data from the 4 corresponding control experiments are pooled for Untreated. For interphase length a minimum of 26 cells was analyzed, and for mitosis length a minimum of 133 cells was analyzed. Statistical tests: Kruskal-Wallis with Dunn’s multiple comparisons tests. **(G- H)** Single cell profiles of PLK4Ctl (G) and PLK4OE (H) Untreated cells. Color coding of mitosis and fates refers to categories defined in Figure 1A, with legend repeated in panel K. **(I)** Averages and SEM of the percentages of mitotic phenotypes (legends in Fig. 1A and panel K). Two independent experiments, statistical test: Fisher’s exact test on the number of *Slight Mis-segregation* events. **(J)** Percentage of multipolar divisions observed in presence or absence of 5nM Paclitaxel. Two independent experiments, statistical test: Fisher’s exact test on the number of multipolar divisions. **(K)** Legends for panels G-H and I, as defined in Fig. 1A.

**Fig. S2.**
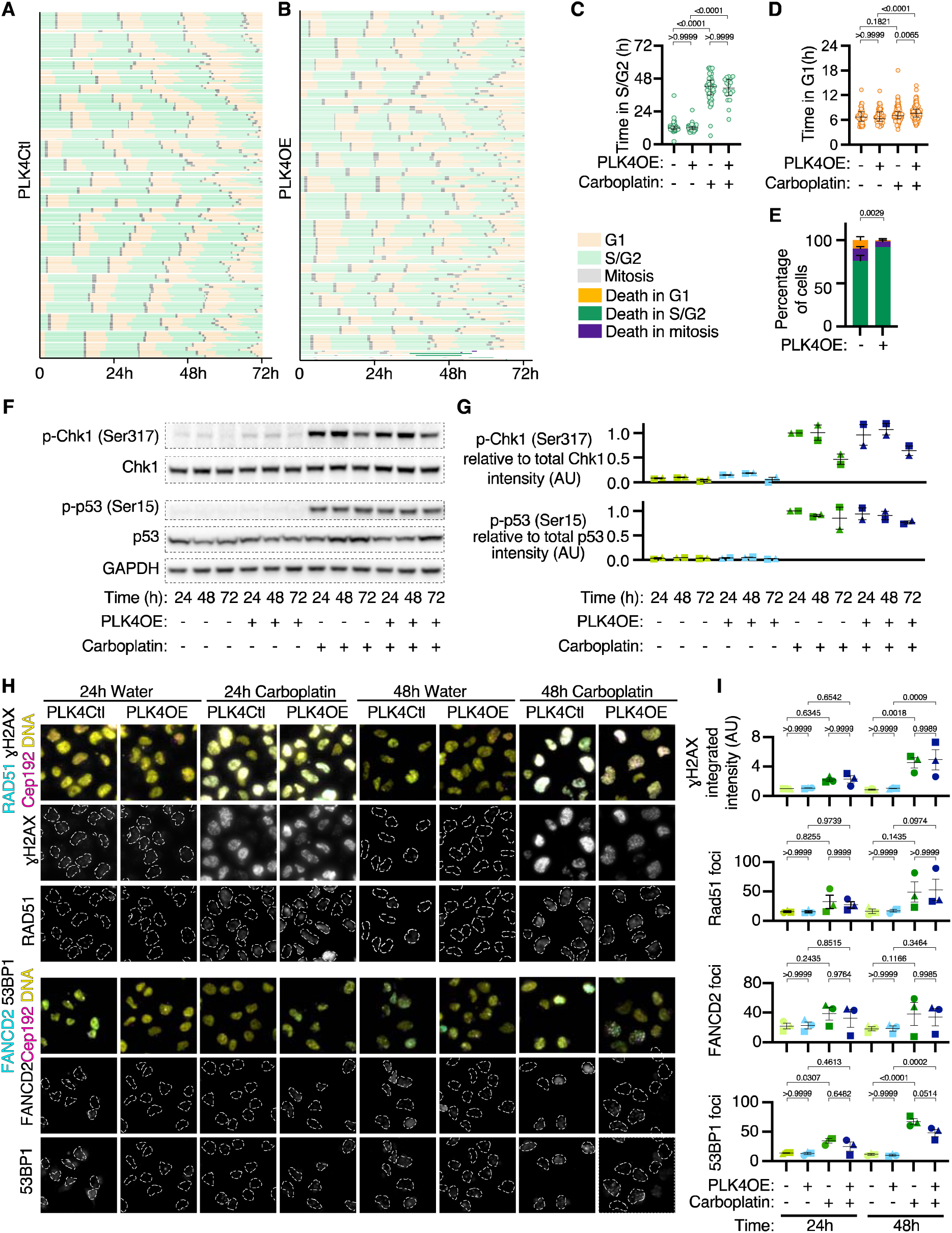
**(A-B)** Single cell profiles of FUCCI PLK4Ctl (A) and PLK4OE (B) untreated cells. See panel B for color-coded legends of cell cycle phase and cell fate. **(C-D)** Scatter dot plot graph of S/G2 (C) and G1 (D) phase lengths in the second generation, with median and interquartile range. Two independent experiments with a minimum of 18 times analyzed per category. Statistical tests: Kruskal-Wallis with Dunn’s multiple comparisons tests. **(E)** Bar graphs showing the averages and SEM of the percentages of cell death events occurring in the indicated cell-cycle phases. Two independent experiments, statistical test: Fisher’s exact test on number of death events occurring in S/G2. **(F)** Representative images of Western Blot analysis of phosphorylated Chk1 and p53 **(G)** Graph showing the average and SEM of phosphorylated protein relative to total protein levels, normalized to the levels detected in PLK4OE cells treated with Carboplatin for 24h, from 2 independent experiments. **(H)** Representative images of cells stained with DAPI an antibodies against FANCD2 (Cyan), 53BP1 (gray) and CEP192 (Magenta). Grayscale images of RAD51 and γH2AX are shown. **(I)** Dot-plot representing integrated nuclear γH2AX fluorescence intensity per cell, or numbers of Rad51, FANCD2 or 53BP1 foci per cells. Average and SEM of the averages obtained from 3 independent experiments, each quantifying a minimum of 94 cells per condition. Values are normalized to the average of untreated PLK4Ctl cells at 24h. Statistical test: ANOVA with Sidak’s multiple comparison tests.

**Fig. S3.**
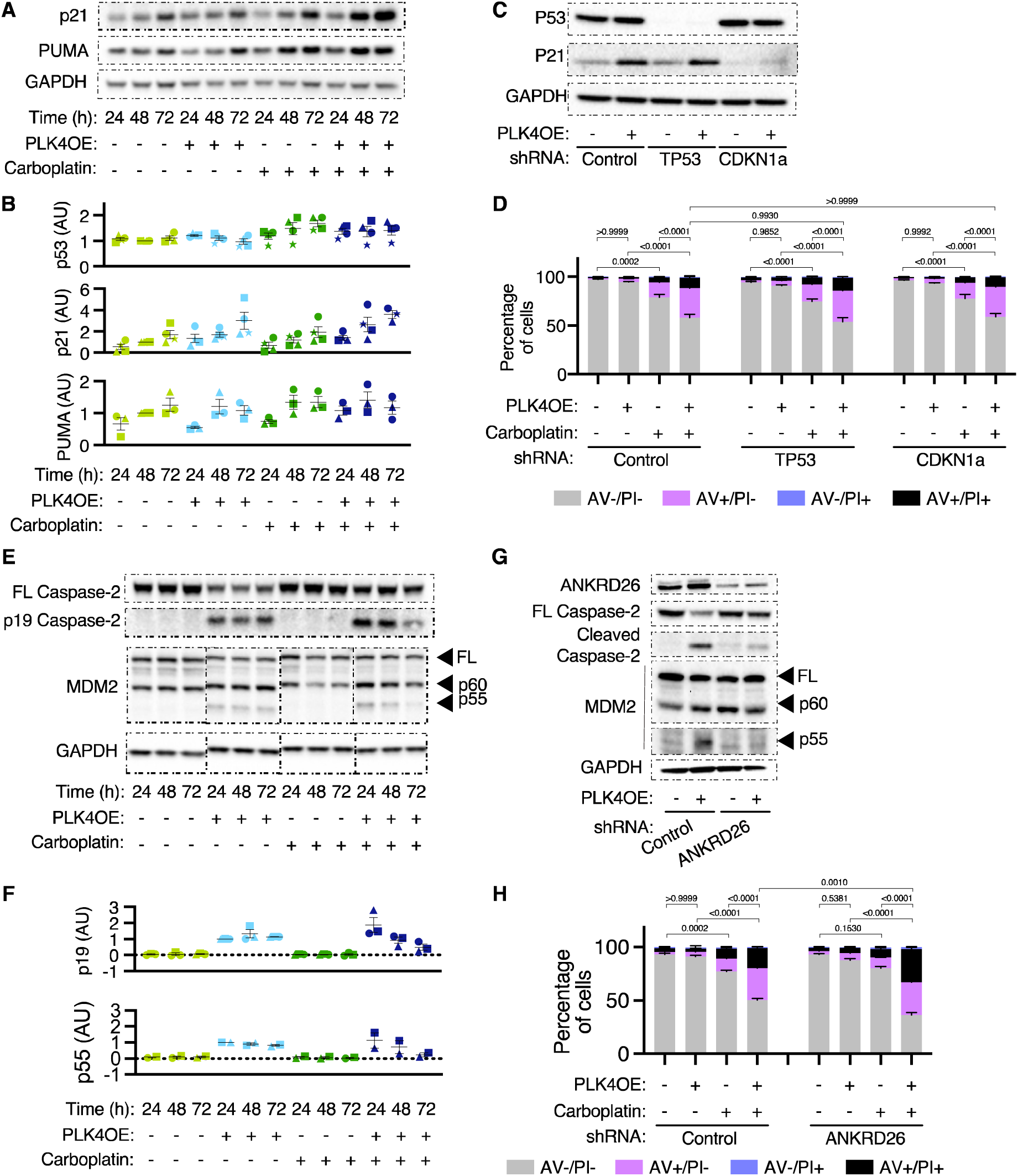
**(A)** Representative images of Western Blot analysis of p21, and PUMA. **(B)** Graph showing the average and SEM of protein levels from 4 (p53 and p21) or 3 (PUMA) independent experiments, normalized to levels measured in untreated PLK4Ctl cells at 48h. **(C and G)** Representative images of Western blot analysis of indicated shRNA cell lines. **(D and H)** Bar graphs showing the average and SEM of the percentage of cells in specified Annexin V-APC/PI gates analyzed by flow cytometry. (D) 6 replicates obtained from 4 independent experiments, with a minimum of 10000 cells analyzed per condition and replicate. (H) 4 replicates obtained from 2 independent experiments, with a minimum of 10000 cells analyzed per condition and replicate. Statistical test: comparison of the percentage of Annexin V positive cells, using ANOVA with Sidak’s multiple comparison test. Representative cytometry profiles can be found in the supplementary materials. **(E)** Representative images of Western Blot analysis of Caspase2 and MDM2 cleavage. **(F)** Average and SEM of protein levels from 3 (p19 Caspase2) or 2 (p55 MDM2) independent experiments, normalized to levels measured in untreated PLK4OE cells at 24h.

**Fig. S4.**
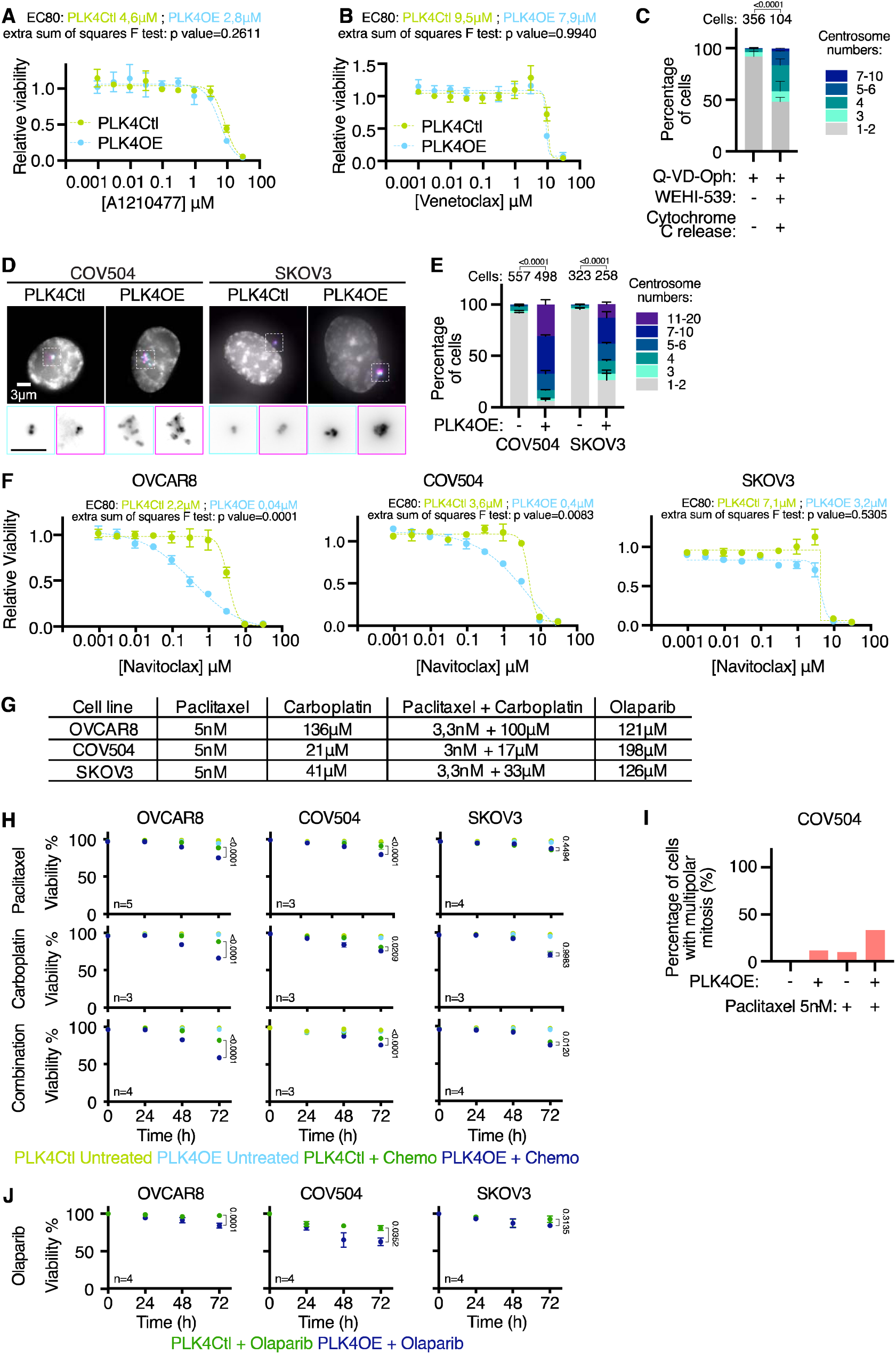
**(A-B)** Dose-response of PLK4Ctl and PLK4OE cells to A1210477 **(A)** and Venetoclax **(B)**, normalized to their respective untreated conditions, obtained from MTT viability assays. Mean and SEM of 2 independent experiments each obtained from averaging 3 technical replicates. **(C)** Bar graphs showing the average and SEM of the percentage of cells with the indicated number of centrosomes (CEP192 dots colocalizing with Pericentrin). 2 independent experiments, statistical test: Fisher’s exact test comparing the number of cells with more than 2 centrosomes. Numbers on the top of each graph represent the number of cells analyzed per condition. **(D)** Representative images of COV504 and SKOV3 cells stained with DAPI (gray) and antibodies against CEP192 (Cyan) and Pericentrin (Magenta). **(E)** Bar graphs showing the average and SEM of the percentages of cells with the indicated number of centrosomes (CEP192 dots colocalizing with Pericentrin). 3 independent experiments, statistical tests: Fisher’s exact tests comparing the number of cells with more than 2 centrosomes. Numbers on the top of each graph represent the number of cells analyzed per condition. **(F)** Dose-response of PLK4Ctl and PLK4OE OVCAR8 (Left), COV504 (Middle) and SKOV3 (Right) to Navitoclax, normalized to their respective untreated conditions, obtained from MTT viability assays. Mean and SEM of 2-3 independent experiments each obtained from averaging 3 technical replicates. **(G)** Table summarizing the IC50s and drug concentrations used in combinations, determined via MTT dose-response viability assays, for PLK4Ctl OVCAR8, COV504 and SKOV3. **(H)** Viability (% of Trypan Blue negative cells) counted for PLK4Ctl and PLK4OE OVCAR8 (Left), COV504 (Middle) and SKOV3 (Right), in response to indicated chemotherapies using concentrations indicated in panel G for 72h. Average and SEM of the number of independent experiments indicated, each obtained from averaging 3 technical replicates. Statistical test: two-way ANOVA with Sidak’s multiple comparison test. **(I)** Percentage of multipolar divisions observed in COV504 in response to 5nM Paclitaxel exposure. **(J)** Viability (% of Trypan Blue negative cells) counted for PLK4Ctl and PLK4OE OVCAR8 (Left), COV504 (Middle) and SKOV3 (Right), in response to Olaparib using concentrations indicated in panel G for 72h. Average and SEM of 4 independent experiments, each obtained from averaging 3 technical replicates. Statistical test: two-way ANOVA with Sidak’s multiple comparison test.

**Fig. S5.**
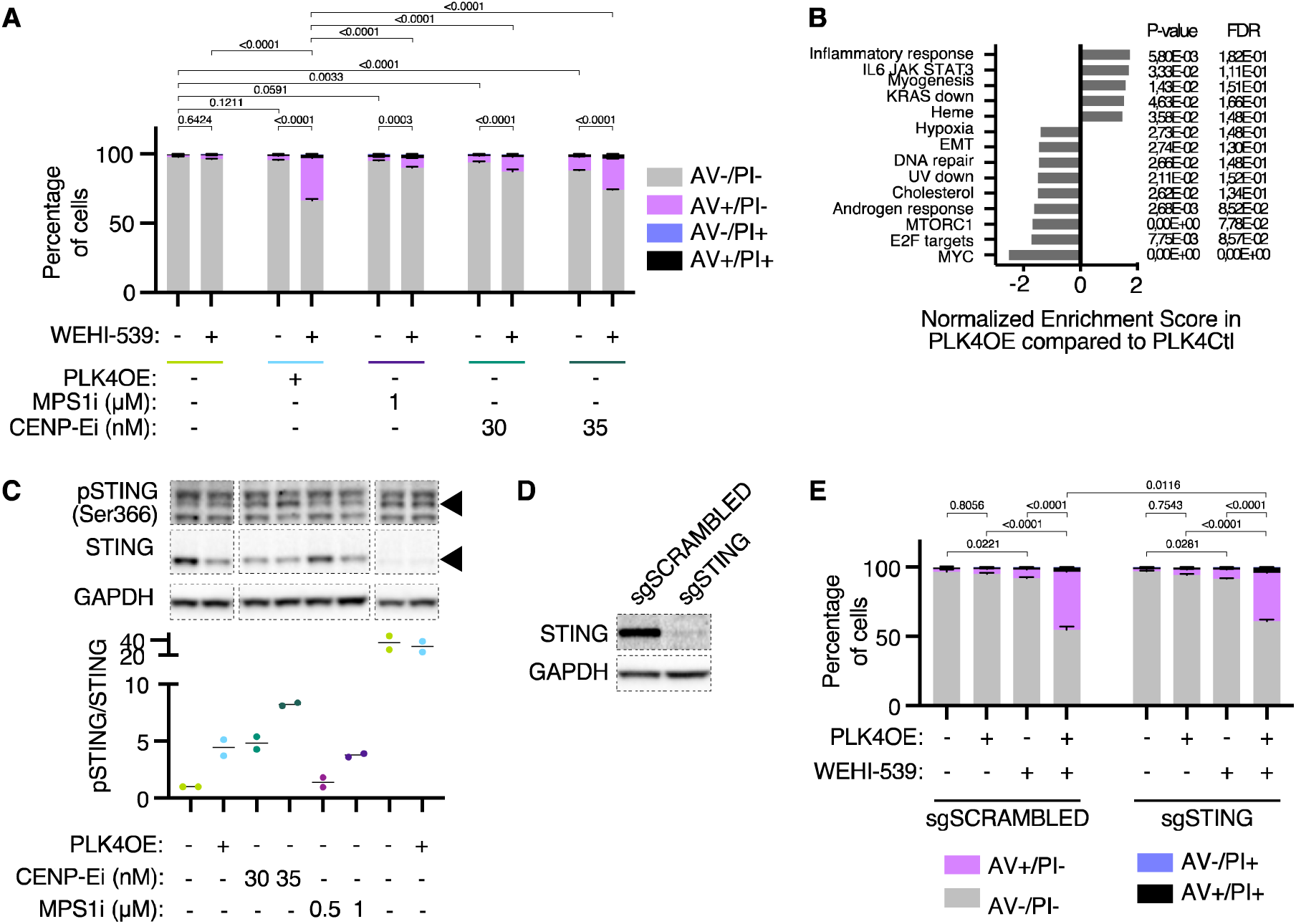
**(A)** Bar graphs showing the average and SEM of the percentage of cells in specified Annexin V-APC/PI gates analyzed by flow cytometry. 3 replicates obtained from 2 independent experiments, with a minimum of 20000 cells analyzed per condition and replicate. Statistical test: comparison of the percentage of Annexin V positive cells, using ANOVA with Sidak’s multiple comparison test. Representative cytometry profiles can be found in the supplementary materials. **(B)** GSEA Hallmarks with |normalized enrichment score >1,5 and p-value < 0,05, from differential RNA expression analysis of PLK4OE cells compared to PLK4Ctl. **(C)** Western-Blot analysis of STING phosphorylation after 72hr of indicated drug treatments with quantification of pSTING relative to total STING. Average and SEM of 2 independent experiments, normalized to untreated PLK4Ctl. HT-DNA transfection was performed 24h before cell collection. **(D)** Western blot analysis of indicated bulk LentiCRISPR cell lines. **(E)** Bar graphs showing the average and SEM of the percentage of cells in specified Annexin V-APC/PI gates analyzed by flow cytometry. 4 replicates obtained from 2 independent experiments with a minimum of 15000 cells analyzed per condition and replicate. Statistical test: comparison of the percentage of Annexin V positive cells using ANOVA with Sidak’s multiple comparison test. Representative cytometry profiles can be found in the supplementary materials.

**Fig. S6.**
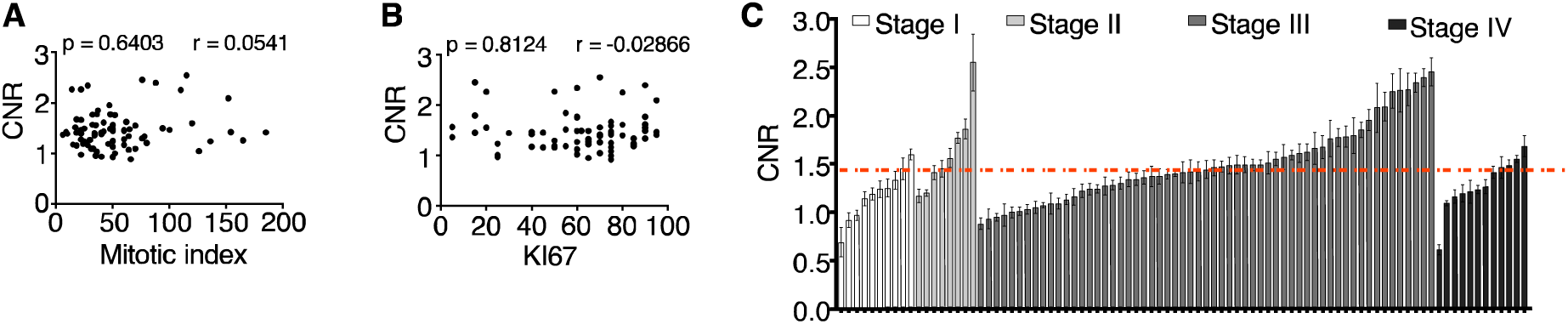
**(A-B)** Distribution of the Mitotic Index (A) and of the percentage of Ki67 positive cells (B) as a function of CNR. Statistical test: Spearman correlation. **(C)** Average and SEM of CNR per patient classified depending on FIGO stage.

### Supplementary Tables

**Table S1.**
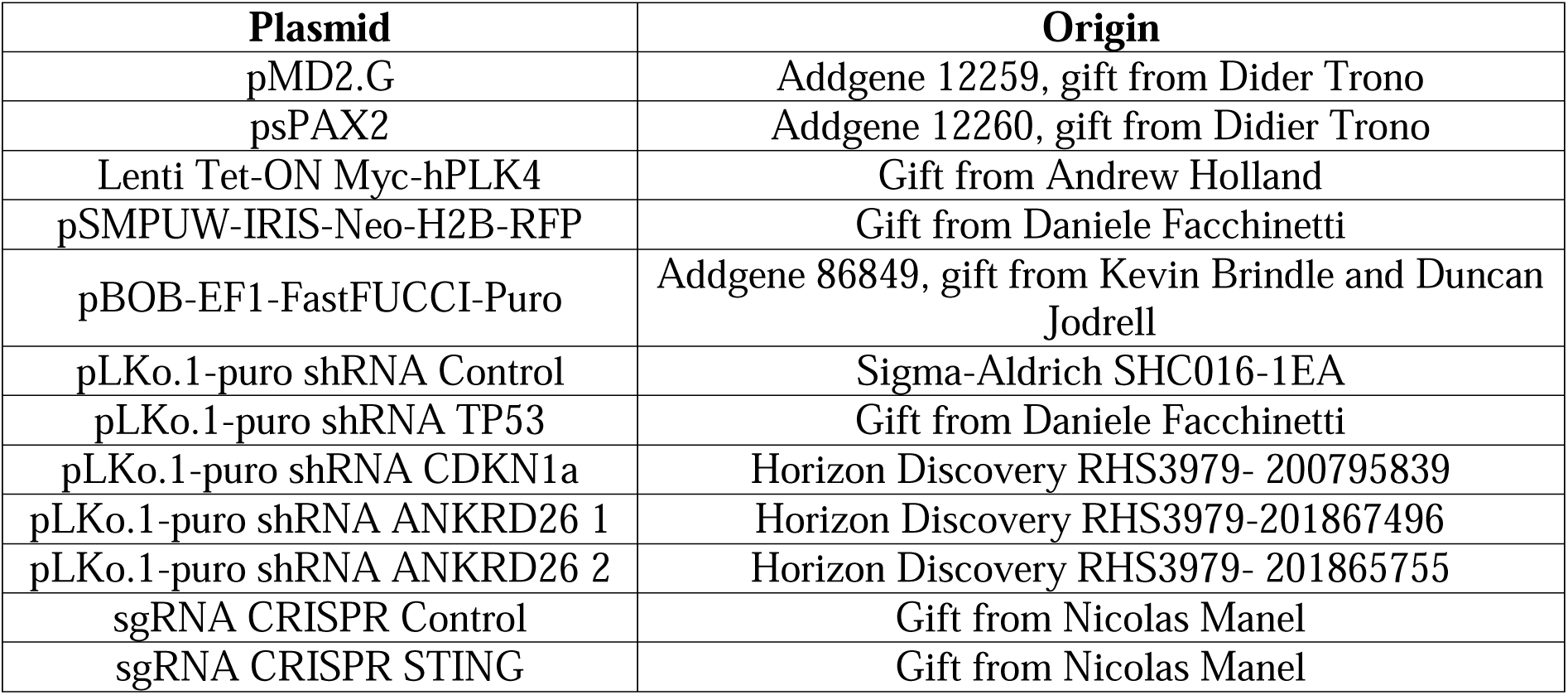
List of plasmids.

**Table S2.**
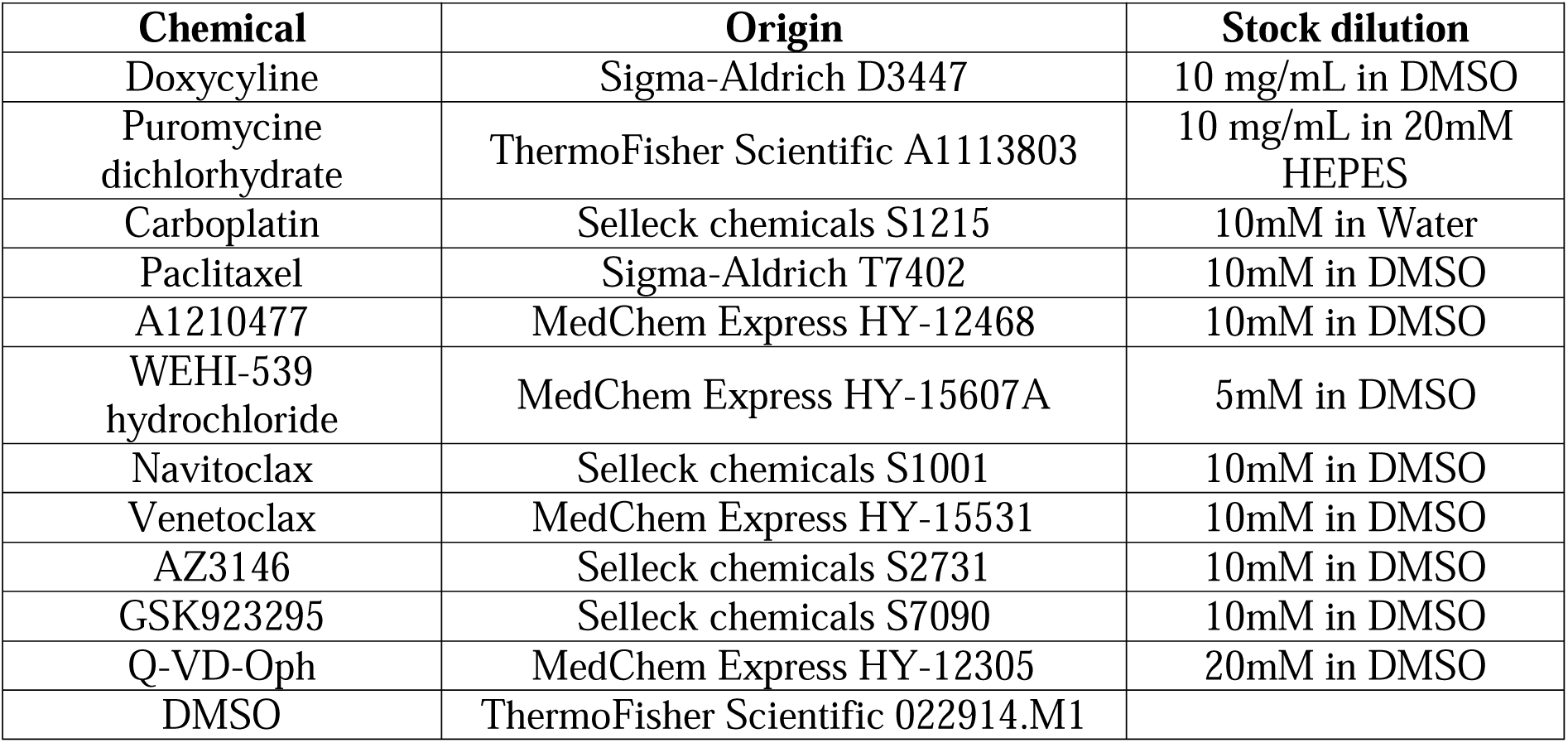
List of drugs.

| **Chemical** | **Origin** | **Stock dilution** |
| --- | --- | --- |
| Doxycycline | Sigma-Aldrich D3447 | 10 mg/mL in DMSO |
| Puromycine dichlorhydrate | ThermoFisher Scientific A1113803 | 10 mg/mL in 20mM HEPES |
| Carboplatin | Selleck chemicals S1215 | 10mM in Water |
| Paclitaxel | Sigma-Aldrich T7402 | 10mM in DMSO |
| A1210477 | MedChem Express HY-12468 | 10mM in DMSO |
| WEHI-539 hydrochloride | MedChem Express HY-15607A | 5mM in DMSO |
| Navitoclax | Selleck chemicals S1001 | 10mM in DMSO |
| Venetoclax | MedChem Express HY-15531 | 10mM in DMSO |
| AZ3146 | Selleck chemicals S2731 | 10mM in DMSO |
| GSK923295 | Selleck chemicals S7090 | 10mM in DMSO |
| Q-VD-Oph | MedChem Express HY-12305 | 20mM in DMSO |
| DMSO | ThermoFisher Scientific 022914.M1 |  |

**Table S3.**
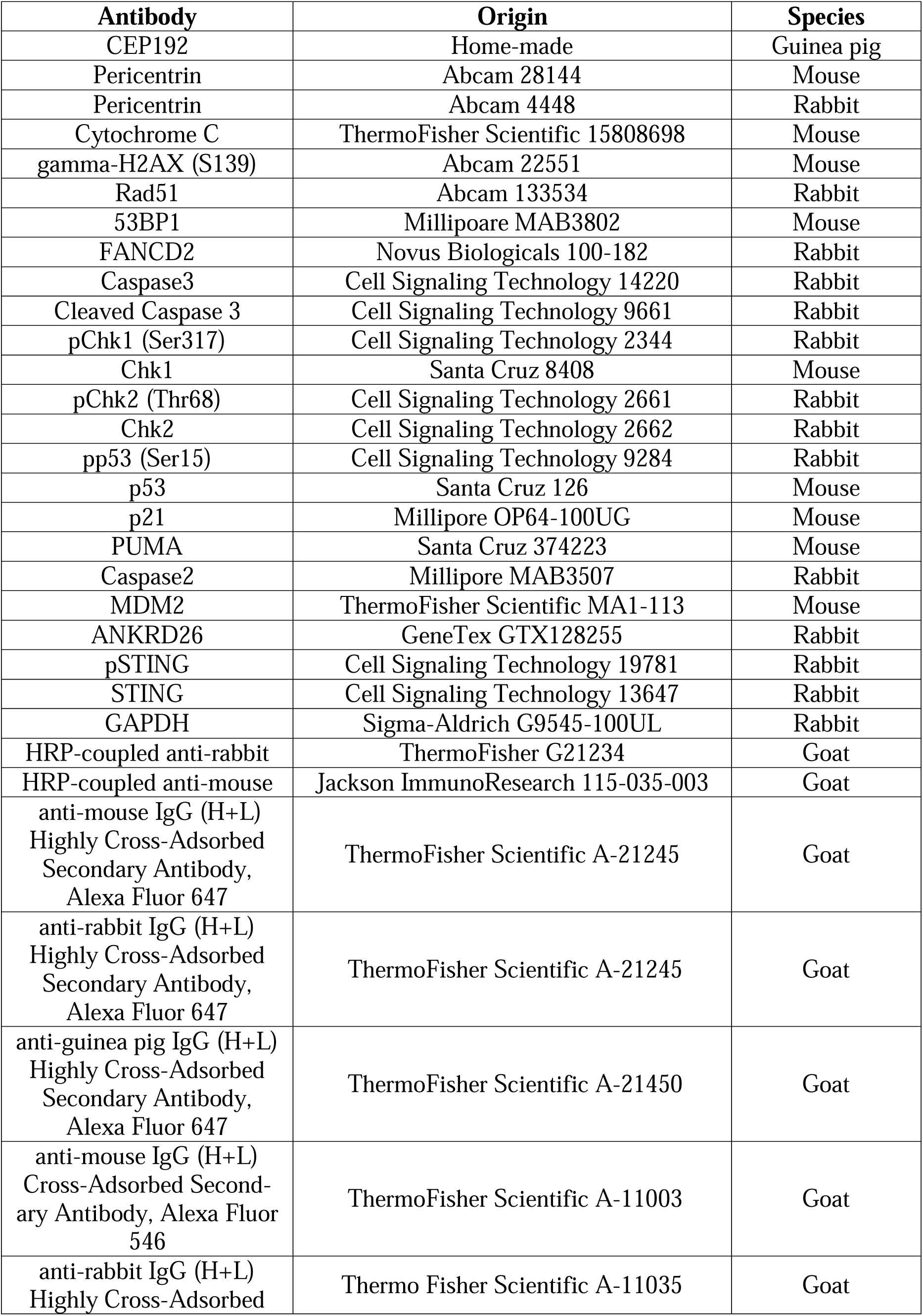
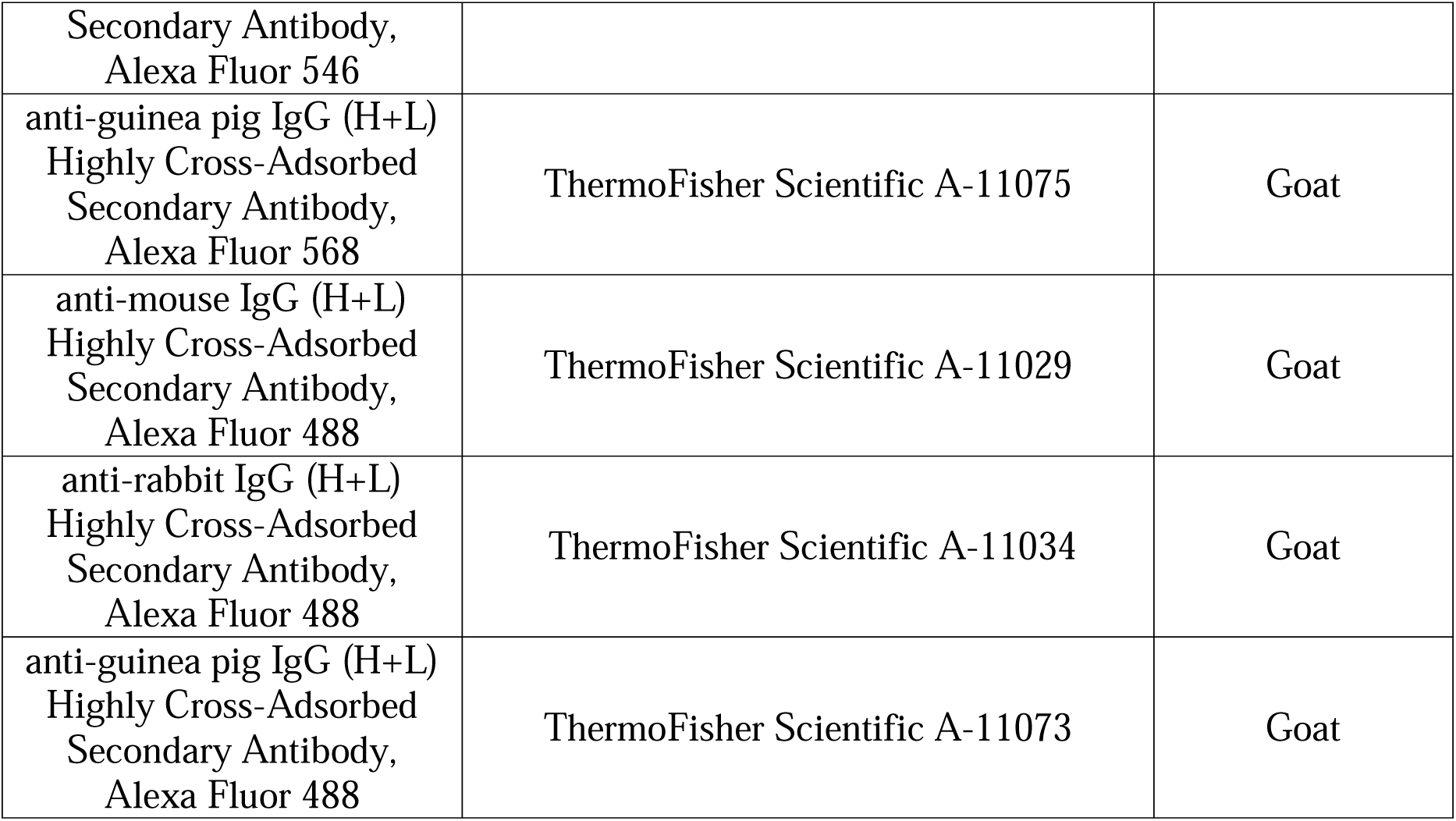
List of antibodies.

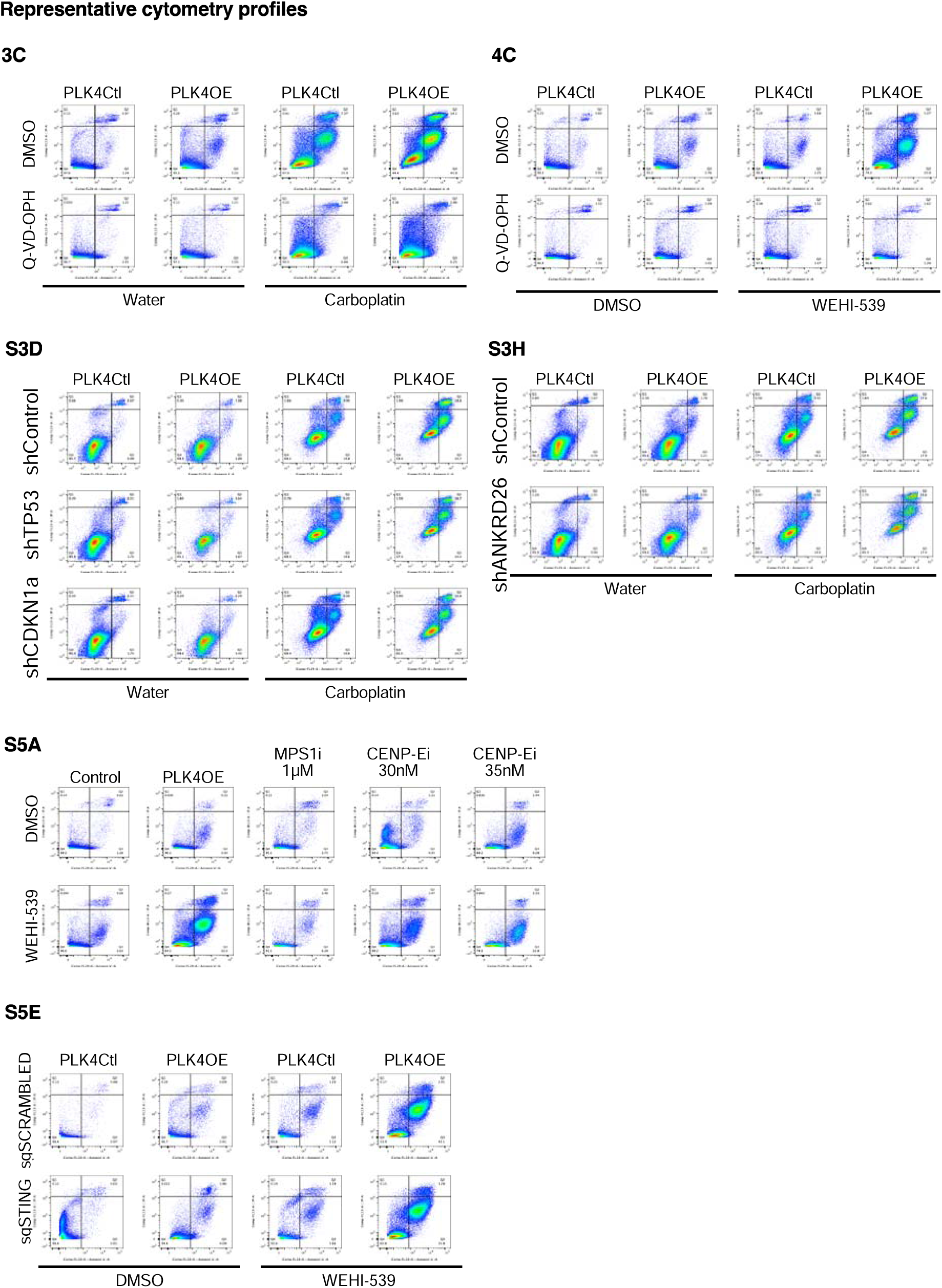

